# Intestinal epithelial *Casd1* influences mucus sialic acid *O-*acetylation and tissue damage susceptibility toward large-intestinal mucosal insults

**DOI:** 10.64898/2026.02.07.702670

**Authors:** Simin Jafaripour, Mackenzie Melvin, Megan Turluk, Emily Howard, Noah Fancy, Hongyuan Zhang, Jackline D.W. Irungu, Brian R. Wasik, Daniel Chou, Benjamin Bigiremana, Nitin, Caixia Ma, Qiaochu Liang, Negin Kazemian, Sepideh Pakpour, Colin R. Parrish, Bruce A. Vallance, Wesley F. Zandberg, Kirk S. B. Bergstrom

## Abstract

The intestinal mucus network, primarily composed of *O*-glycosylated MUC2 mucin polymers, is essential for protecting the gastrointestinal tract from microbial threats. Sialic acid (Sia), a terminal monosaccharide on complex *O*-glycans, plays a key role in maintaining mucus integrity and is frequently modified by Casd1-dependent *O*-acetylation (*O*Ac). Despite its prevalence, the biological significance of sialic acid *O*Ac (*O*Ac-Sia) modifications in human and murine mucus remains unclear. We hypothesized that *O*Ac-Sia variants on mucus interact with the microbiota and are required for optimal mucus barrier function and host-microbe homeostasis in the colon. To test this, we profiled *O*Ac-Sia on human and mouse MUC2 in situ using viral-derived probes with bacterial FISH and confocal microscopy; generated intestinal epithelial cell (IEC)-specific Casd1 null mice (IEC *Casd1^-/-^*); performed sialylomic and *O*-glycomic HPLC-MS analyses; assessed microbial communities by 16S rRNA sequencing with quantitative microbial profiling (QMP); and evaluated disease susceptibility using DSS colitis and *Citrobacter rodentium* infection models. Results revealed that both human and murine mucins are extensively *O*-acetylated and interact with the microbiota, suggesting biological relevance. IEC *Casd1^-/-^* mice were viable and displayed a complete loss of mucin *O*Ac-Sia, indicating Casd1 is the sole contributor to *O*Ac-status. Unexpectedly, mucus function was intact in IEC *Casd1^-/-^* mice, with no difference in structure or quality vs. WT co-housed littermates.16S rRNA analysis showed a modest but significant sex-specific reduction of microbial loads in male IEC *Casd1^-/-^*mice, and a clear trend toward reduced *Turicibacter* spp. vs. WT mice in both male and females, without impacting overall short-chain fatty acid (SCFA) production. DSS treatment led to more severe and extensive tissue damage in IEC *Casd1^-/-^* mice. *C. rodentium* infection led to increased damage in the cecum and distal colon of IEC *Casd1^-/-^* mice without affecting pathogen load, suggesting that *O*Ac-Sia status has a role in tolerance defense. These findings establish intestinal epithelial Sia *O*-acetylation as a component dispensable for mucus and host-microbe homeostasis at baseline, but important in limiting damage to mucosal inflammatory insults.

## INTRODUCTION

Mucus barrier function is essential for intestinal homeostasis. Mucus is a highly glycosylated polymeric network that forms the interface between microbes and humans in the gut^1,2^. Mucus serves two major functions: being a niche and food source for a subset of microbes in the gut, and a barrier to limit microbial intrusion into the gut wall^3^. Its importance is established in animal models, showing loss of MUC2 predisposes to chronic colitis^4–6^, cancer^7^, and enteric infection^8,9^. Recent studies have shown this interface is mobile; that is, it adheres to the feces in both mice and humans^1,10,11^. In mice, the barrier function of mucus is composed of two sub-layers, a major layer derived from the proximal half of the colon (the “b1” layer), and a highly sulfated minor layer coming from the distal colonic goblet cells (the “b2” layer), characterized by high affinity for the lectin *Maakia amurensis* Lectin (MAL) II^10^. Both layers contribute to barrier function and homeostasis^10^.

Mucus is made up of the mucin MUC2 and is mainly comprised of oligosaccharides attached to the MUC2 backbone. These glycans are primarily *O*-linked (*O*-GalNAc), while ∼10% are N-linked^12^. The MUC2 glycans dictate most function of mucus. N-glycans control MUC2 folding and ER stress, as loss of N-glycan enzyme Mpi disrupts mucus biosynthesis and folding in the ER and prevents its establishment in the colon^13^. Conversely, *O*-glycans regulate its function in the lumen, as loss of complex glycans destabilizes the mucus leading to chronic microbiota-dependent colitis and colitis-associated cancer^14,15^.

Recent work has focused on sialic acid (Sia), an essential capping monosaccharide on mucus that influences mucus-microbe interactions in the gut^16–18^. On mucins, Sia is typically linked to N- and *O*-glycans via α2,3 and/or α2,6 to subterminal Gal and/or GalNAc residues on non-reducing ends^19^. Sia is a nine-carbon sugar with a six-membered ring and a glycerol-like side chain via carbons (C)7–9, and is negatively charged due to C1 being modified by a carboxyl group that is deprotonated at physiologic pH^20^. Sialic acid (Sia) plays central protective roles in mucus barrier integrity preventing dybisosis, with loss-of-function mutations in ST6GalNAc1 ^17^and ST6GalNAc6 ^18^ impairing mucus structure and increasing susceptibility to colitis and mucosal injury. We have shown that global loss of Sia via disruption of the CMP-Sia transporter Slc35a1 drives major microbiota alterations, mucus defects in the b1 barrier layer, and spontaneous microbiota-dependent inflammation and tumorigenesis^16^. Conversely, Sia can also fuel opportunistic pathogens^21^, enhance virulence^22^, and promote proinflammatory bacterial outgrowth^23^. These studies highlight the need for tightly regulated sialylation to maintain intestinal homeostasis and protection from infection.

The numerous hydroxyl (-OH) groups are sites of extensive modification by various functional groups, the best studied of which are *O*-acetyl groups^20^. These are found either singly on Sia (e.g., Neu5,9Ac) or in various combinations (Neu5,7,9Ac; Neu5,7,8Ac), typically on C7–9, but may also include the C4 position^24^. The *O*Ac groups influence the functional biology of Sia^25^. For example, the *O*Ac motifs on Sia can serve as binding sites for viral hemagglutinins^26,27^, can inhibit neuraminidase activities (which influences how long Sia can remain on glycoconjugates^28,29^), and can influence lectin binding, with implications for Siglecs and immune functions^25,30,31^. Biosynthesis of *O*Ac variants is governed by sialate *O*-acetyltransferases, of which at least two have been discovered in the human genome. The best studied is CAS1 domain-containing protein 1 (CASD1), which has preference for activated nucleotide sugar CMP-Neu5Ac prior to its translocation into the Golgi via Slc35a1 and subsequent incorporation into glycans via sialyltransferases (STs) in the Golgi^32,33^. CASD1/Casd1 can modify Sia at both 7- and 9-positions^32^, and this reaction can be followed by migration of *O*Ac to different C residues either spontaneously or via a possible enzyme^31^, or reversed through actions of sialated *O*-acetyl esterase (SIAE)^27^. CASD1 is widely expressed throughout the body^25,34^ (Human Protein Atlas), suggesting cell- and tissue-specific roles. Recently, Varki and Eckmann and colleagues (2025) demonstrated that global Casd1 loss in mice ablates Sia *O*-acetylation in tissues, and predisposes to acute colitis caused by the colitogenic cytotoxin dextran sodium sulfate (DSS)^35^. This finding highlights the importance of *O*Ac-Sia; however, the cell types responsible remain to be identified. The second enzyme is Neurexophilin PC-domain containing protein-1 (NXPE1), which has a similar active site as CASD1 and is highly expressed in human colonic epithelial cells, and mainly targets Sia at the C9 position^36,37^. Notably, a rare variant of NXPE1 (NXPE1G533R) was identified in genome-wide association studies, and was associated with protection from Ulcerative Colitis (UC)^37^. Collectively, these studies suggest a complex relationship between OAc status and disease susceptibility.

Despite the extensive *O*Ac modifications on colon mucus^38,39^, the role of *O*Ac-Sia in regulating Sia and mucus biology is unclear. Lewis and colleagues have suggested a role in regulating accessibility of pathogenic *E. coli* to Sia for energy and growth in vitro^28^, suggesting a role in regulating mucosal ecology. This is supported by microbiota profiling in *Casd1*^-/-^colons^35^. Functional characterization of the rare UC-protective NXPE1G533R variant ^37^ revealed this to be a loss-of-function mutation suggesting *O*-acetylation of mucus may not be beneficial in UC^37^. In contrast, de-*O*-acetylation via inflammation-dependent expression of SIAE has been suggested to render the sialome more prone to hydrolysis, compromising the mucus barrier and predisposing to colitis^40^. These conflicting conclusions can only be resolved by comparing mucus function with and without *O*Ac-Sia on mucus.

We now report the first formal investigation into the cells responsible for the biological function of *O*Ac-Sia in the mammalian colon and its relevance to mucus function. By generating a model of gut-epithelium-restricted ablation of *O*Ac-Sia, we provide evidence that epithelial-intrinsic Sia *O*-acetylation is not required for baseline mucus stability and overall function or homeostasis. However, *O*Ac-Sia does have a subtle sex-specific impact on gut microbial ecology as shown through relative and quantitative microbial profiling. *O*Ac-Sia also has a protective effect against mucosal damage caused by acute toxins such as DSS, as well as enteric infection caused by *C. rodentium,* but not bacterial burden. These studies indicate that *O*-acetyl modification of Sia is a mechanism to limit damage and inflammation caused by chemical or microbial toxins, but have no detectable roles in directly regulating mucus barrier function at baseline.

## RESULTS

### *O*-acetylated Sia are associated with the microbiota in humans and mice

Several studies have profiled the diversity of Sias in human colonic mucus^39,41^; however, their spatial expression in relation to fecal mucus is unknown. To address this in healthy individuals, we leveraged recent findings on the utility of human feces to probe mucus non-invasively *in situ*^1^. Using Carnoy’s-fixed paraffin-embedded (CFPE) human fecal sections, we probed for different *O*-acetylated sialic acid variants using recombinant human IgG Fc–tagged probes targeting 9-*O*Ac (Porcine Torovirus P4 esterase, PToV HE), 7,9-*O*Ac (Bovine Coronavirus Mebus strain esterase, BCoV-M HE), and 4-*O*Ac-Sia (Mouse hepatitis virus S strain *O*-acetylesterase, MHV-S HE) derivatives^42^. We found that the human MUC2 barrier layer stained abundantly for 9- and 7,9-di-*O*Ac, clearly marking the mucus encapsulation at the edge of the fecal mass (**Fig. 1A**). Interestingly, we also observed a clear signal for 4-*O*Ac, although this was concentrated at the edge of the pellet above the mucus, with its signal radiating further into the fecal mass. In all cases, target specificity was confirmed using negative control probes (esterase-positive, removing the *O*Ac group upon binding, or use of a non-binding mutant) (**Fig. 1A**).

**Figure 1.**
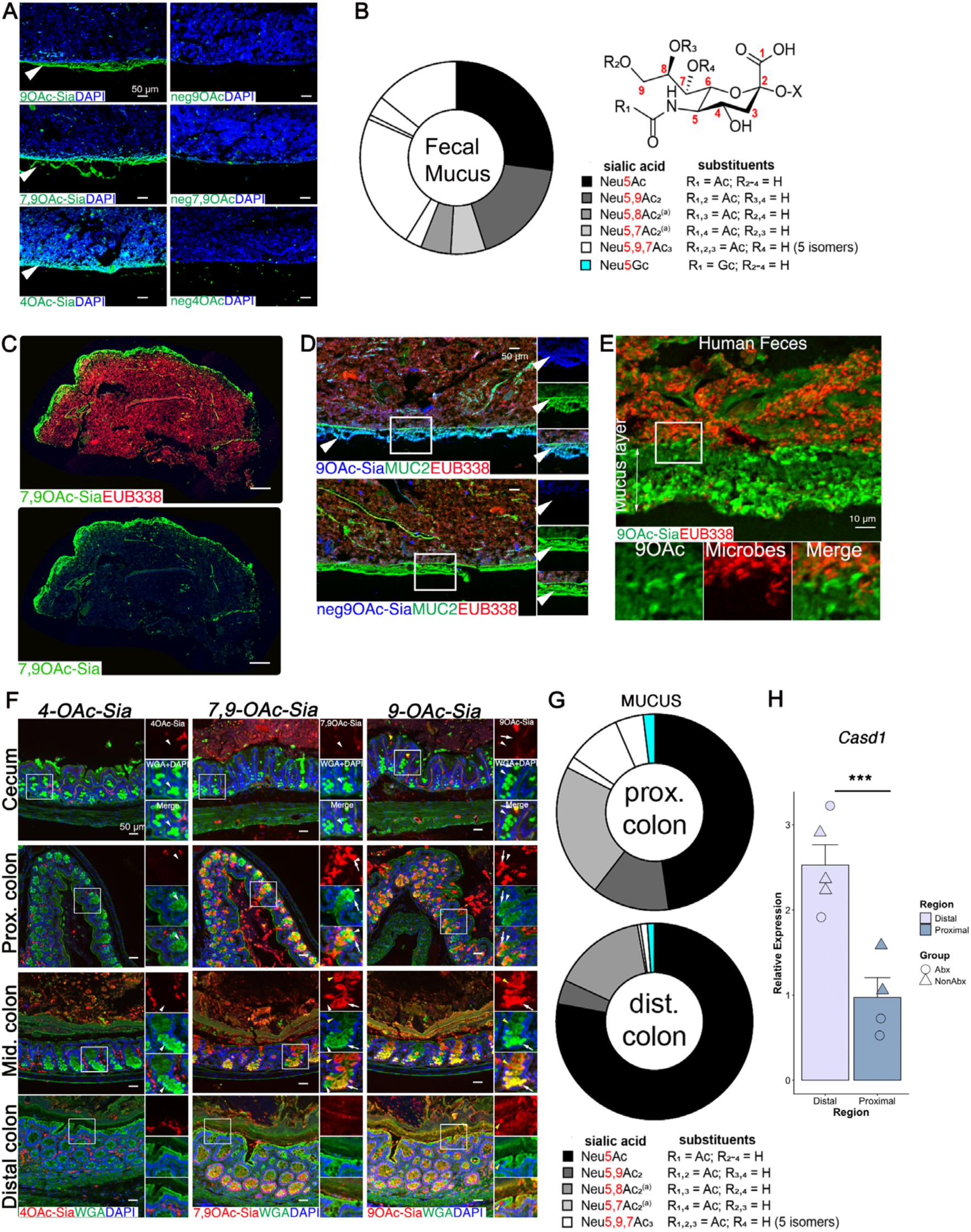
Spatial and relative distribution of *O*-acetylated Sia isoforms in human and murine colonic mucus. **A**. Epifluorescent imaging of *O*Ac-Sia variants on human fecal mucus. **B**. Sialylomics analysis of human fecal MUC2. **C**. Tiled image of *O*Ac-Probe:FISH-stained CFPE healthy human feces showing overall distribution of 7,9-di-*O*Ac Sia (green) on fecal human MUC2 in relation to the microbiota (red). **D**. High-magnification image of *O*Ac-Probe:FISH-stained CFPE healthy human feces showing 9-*O-*acetylation is associated with barrier function. **E**. Confocal imaging of 9-*O*Ac-Probe:FISH on CFPE healthy human feces showing intimate interactions of 9-*O*-acetylated Sia on human mucus with the gut microbiota. Image is representative of four healthy colons. **F**. Epifluorescent imaging of 9-, 7,9-, and 4-*O*-acetylated Sia variants in tissues and mucus. **G**. Gene expression analysis (mean ± SEM) of *Casd1* in mouse epithelium derived from proximal and distal colon. Data points indicate individual mice. **H**. Sialylomics of mucus extracted from proximal and distal colons of WT mice showing region-specific relative abundances of different Sia isoforms.

To achieve higher resolution and determine the proportion of Sia modified by *O-*acetylation, we extracted MUC2 from human fecal material using established protocols^1^, derivatized the Sias, and performed sialylomics using HPLC–MS. We found that 75% of MUC2 was *O*-acetylated, with varying levels of mono-, di-, and tri-*O*-acetylated Sias, and 9-*O*Ac dominating among the *O*-acetylated variants (**Fig. 1B**). To determine interaction with the microbiota, we co-labeled BCoV HE (7,9-*O*Ac–targeting)–stained fecal sections with the bacterial probe EUB338-TexRd and imaged entire sections. We found that 7,9-*O*Ac–sialylated mucus surrounded the microbial mass (**Fig. 1C**). Similar results were seen with 9-*O*Ac Sia, although it was primarily associated with the barrier layer (**Fig. 1D**). High-magnification confocal imaging confirmed the intimate interaction of the microbiota with *O*Ac-Sia on human colonic mucus (**Fig. 1E**). These studies show that Sia on human MUC2 is richly *O*-acetylated and interacts with the microbiota.

We next determined whether similar phenotypes were observed in mice by analyzing CFPE sections throughout the intestinal tract. We found that distal colon goblet cells and the fecal mucus barrier stained strongly positive for PToV HE and BCoV HE (9- and 7,9-*O*Ac Sia, respectively), but negative for 4-*O*Ac (**Fig. 1F**). Similar staining patterns were seen in the middle and proximal colon (**Fig. 1F**), suggesting that *O-*acetylation status of MUC2 is conserved between mice and humans. The goblet cells of the cecum and terminal ileum were unexpectedly negative for all *O*Ac variants (**Fig. 1F** and **Fig. S1A**).

Sialylomics of mouse Muc2 from proximal and distal colon showed that, despite higher *Casd1* expression in the distal colon, only ∼10% of Sia was *O*-acetylated, in contrast to the proximal colon where *O*Ac-Sia abundance and diversity were richer, making up ∼40% of Sia (**Fig. 1G**). Recent studies have identified cas-domain–containing protein 1 (*Casd1*) as the major *O*-acetyltransferase governing 9-*O-*acetylation in mice^35^. Notably, *Casd1* is the only known *O*-acetyltransferase in mice, which lack a functional ortholog of the additional human O-Ac transferase *NXPE1*^36,37^. To determine if *O-*acetylation status correlated with *Casd1* in mice, we examined *Casd1* expression in the epithelium of WT mice, finding that *Casd1* mRNA was highly expressed in the mouse epithelium, with higher levels in the distal colon compared to the proximal colon (*p* = 0.0065), in a manner independent of an intact microbiota as determined via broad-spectrum antibiotic treatment (**Fig. 1H**), which is consistent with previous observations in Germ-free vs Conventionally-raised mice^35^. To our knowledge, this is the first detailed in situ analysis of OAc-Sia diversity on mucus in mice and humans, and indicate that diverse *O*-acetylated Sia variants are present on secreted MUC2/Muc2 and readily intermix with the microbiota in a conserved manner between these species.

### Generation of mice with gut-epithelial-restricted deficiency of Sia *O-*acetylation by targeting *Casd1*

To determine the contribution of *Casd1* to overall mucus *O-*acetylation and understand its biologic function in the colon, we used a gene-targeted approach. Although *Casd1^-/-^* mice are available, *Casd1* is widely expressed among tissues, which could limit interpretation of intestinal phenotypes. Therefore, we obtained *Casd1* “floxed” (*Casd1^f/f^*) mice and crossed them with VillinCre transgenic (Tg) mice, which express the loxP (floxed)-targeting Cre recombinase under the control of the epithelial-specific Villin gene promoter, enabling excision of the floxed region only in the intestinal epithelium, leaving it intact in most other cells^43,44^. We define these as intestinal epithelial cell-specific *Casd1^-/-^* or IEC *Casd1^-/-^* (**Fig. 2A**; see also Methods). Unexpectedly, routine genotyping of the *Casd1* tm1c allele (floxed allele) (**Fig. S1B - E**) revealed apparent restoration of WT alleles in ∼6% of offspring from *f/f* × *w/w* crosses, and ∼4% from *f/f* × *f/w* crosses, increasing to ∼50% by the F4 generation, inconsistent with Mendelian inheritance and indicative of locus rearrangement. Structural analysis identified a previously unrecognized variant of the *Casd1*^tm1c^ allele containing two alterations: a 99-bp deletion near the retained Frt site replaced by WT-derived sequence, and a second alteration near the downstream loxP site producing a ∼50-bp larger amplicon based on agarose gel electrophoresis (**Fig. S1D**). The WT allele remained intact, and the modified floxed allele retained functional loxP sites with <100-bp divergence from the canonical allele, suggesting preserved Cre recombination capacity; nevertheless, animals carrying these variants were excluded to re-establish a genetically stable IEC-*Casd1* colony and ensure experimental consistency.

**Figure 2.**
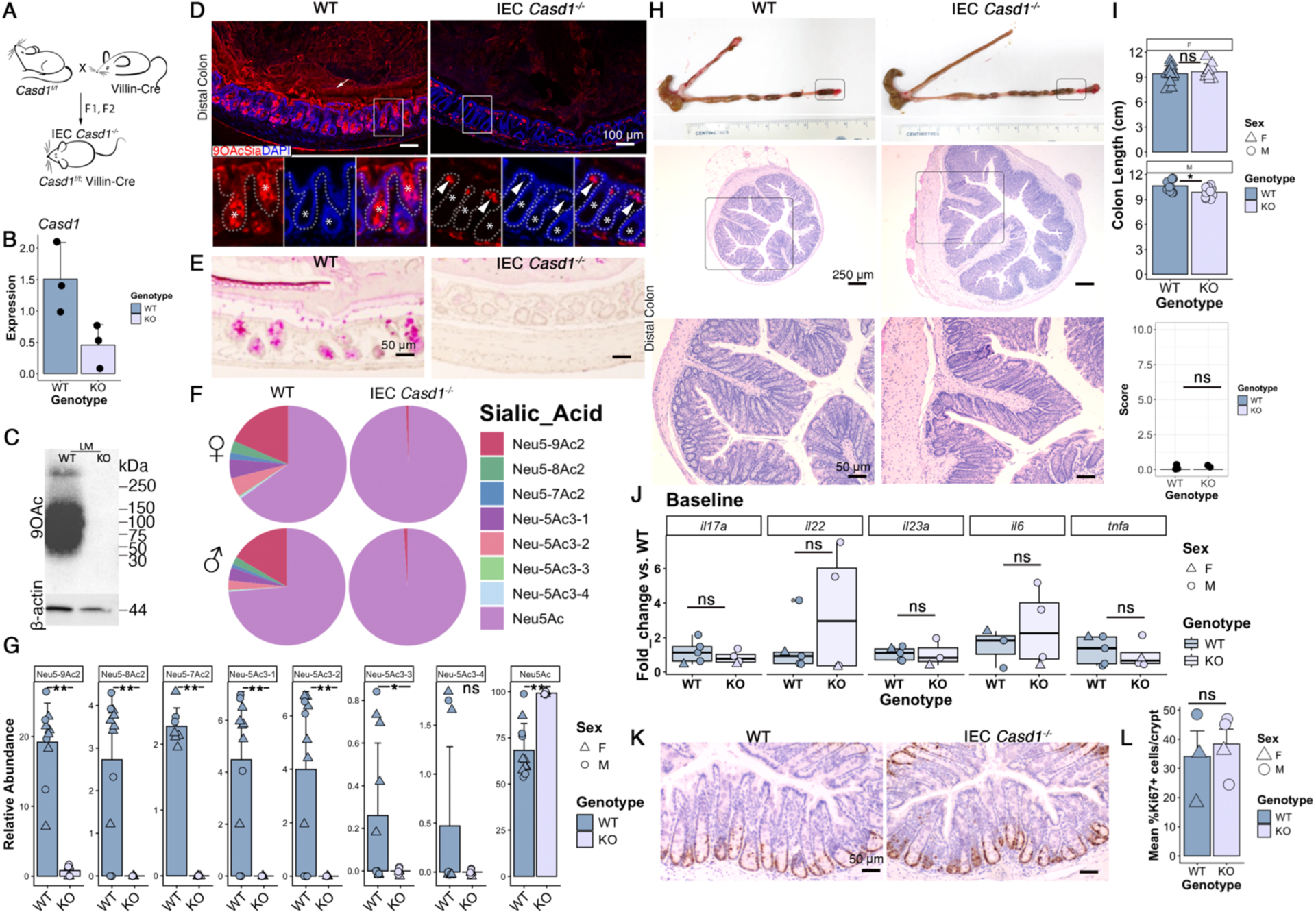
Loss of *Casd1* in epithelium obliterates Sia O-acetylation on colon mucus but does not impair baseline homeostasis. **A**. Schematic of derivation of IEC *Casd1^-/-^* mice in our facility. **B**. Expression of *Casd1* in epithelium of IEC *Casd1^-/-^* vs. WT (*Casd1^f/f^*) mice. **C**. Western blot of 9-*O*-Ac (via PToV4 HE probe) in proteins of the epithelium of WT and IEC *Casd1^-/-^* mice. β-Actin is a housekeeping loading control. **D**. Representative epifluorescent imaging of overall 9-*O-*acetylation *in situ* within CFPE distal colon sections of WT and IEC *Casd1^-/-^* mice. Insets: asterisk = colonic crypt (epithelium), arrow shows retention of 9-*O*Ac signal in the lamina propria (LP). Dashed line separates crypt from LP. **E**. Representative histochemical staining via the PB–KOH–PAS method to visualize 8-*O*-Ac *in situ*. **F**. Pie chart summarizing sialylomics of fecal Muc2 showing impact of *Casd1* on relative abundance of various subsets of *O*-acetylated Sia on mucins. **G**. Bar plots (mean ± SD) overlaid with individual data points showing relative levels of various *O*-Ac subsets between genotypes. (\**p* <0.05, Wilcoxon rank-sum test, FDR-corrected.) **H**. Representative micrograph and histology of mouse (3-month-old) intestinal tissue (n ≥ 8/genotype/sex). Pictures show ileum, cecum, and colon. Histology (H&E) represents the boxed region in the picture. The high-magnification histology is of the boxed region in the above image. **I**. Plot of mean ± SD (overlaid by data points of individual mice) of overall colon length in WT vs. IEC *Casd1^-/-^* mice, stratified by sex. (* = *p* < 0.05, Welch’s t-test.) **J**. Boxplots of gene expression of inflammatory genes in whole distal colon tissues of WT and IEC *Casd1^-/-^* mice. Individual data points are normalized relative expression values within individual mice. **K**. Representative IHC of Ki67 in distal colons. **L**. Right: bar plot (mean ± SEM) overlaid with individual data points of mean percent of Ki67⁺ cells per crypt.

In genetically stable mice with the IEC *Casd1^-/-^* genotype, the epithelial deletion of exon 2 of the *Casd1* gene, along with the resulting loss of its expression and 9-*O*Ac-Sia, in knockout animals was confirmed through the analysis of RNA and protein extracted from IECs compared with their WT littermates (**Fig. 2B & C**, respectively). The efficiency of the knockout system and loss of 9-*O*Ac signal exclusively in the intestinal epithelial cells (IECs) of the IEC *Casd1^-/-^*animals were confirmed *in situ* (**Fig. 2D**). Although 9-*O-*acetylation was virtually gone, we wondered whether there was some compensatory response with other forms of *O*Ac-Sia variants. To first test this, we conducted a histochemical stain of 7,8-*O-*acetylation via Potassium Borate–Potassium Hydroxide–Periodic Acid–Schiff (PB–KOH–PAS) staining, an established protocol for these *O*Ac modifications^45^. We found that while there was a strong signal in WT mice, there was no signal in IEC *Casd1^-/-^* mice (**Fig. 2E**), suggesting that loss of *Casd1* impacts synthesis of other *O*Ac-Sia variants on mucus. This was consistent with a complementary approach using the mildPAS (mPAS) protocol, which labels vicinol diols of reduced non-*O*-acetylated mucins, showing strong labeling in IEC *Casd1^-/-^* mice but faint staining in WT mice (**Fig. S1F**). To confirm this, we extracted colonic mucins from mucus-encapsulated fecal materials, and found that all mono-, di-, or tri-*O*-acetylated Sia species analyzed were absent in the fecal mucus from IEC *Casd1^-/-^* animals of both sexes (**Fig. 2F, G**). These data indicate gut epithelial-intrinsic *Casd1* drives all detectable *O-*acetylation of Sia in mouse colonic mucus.

### IEC *Casd1^-/-^* mice do not exhibit any clinical or histological phenotypes at baseline

In parallel with sialylomics, we performed baseline phenotypic characterization of IEC *Casd1^-/-^*mice vs. WT mice. Both WT and IEC *Casd1^-/-^* littermates, male and female, developed normally with no discernible clinical phenotype. To characterize the intestinal phenotypes of the mice, we first harvested tissues at 3 months of age (**Fig. 2H**). In males, colon length was marginally but significantly greater in WT vs. IEC *Casd1^-/-^* mice (WT: 10.64 ± 0.68 cm, n = 11; KO: 9.87 ± 0.65 cm, n = 10; Welch’s *t*-test, p = 0.016). In females, colon length did not differ between genotypes (WT: 9.42 ± 0.95 cm, n = 17; KO: 9.68 ± 0.94 cm, n = 9; Welch’s *t*-test, *p* = 0.52). (**Fig. 2H and I**). We also processed colon and small intestinal tissues for histology by Hematoxylin and Eosin (H&E) staining. Consistent with the clinical health scores, both groups displayed normal tissue architecture in the distal colon, characterized by well-defined epithelial layers, intact crypt structures, and absence of inflammation (**Fig. 2H**, lower panels). Similar findings were observed in terminal ileum, cecum, and proximal and middle colons (**Fig. S1G**). We also noticed no phenotype in older mice (e.g., 12 months of age, not shown). As a more sensitive detector of baseline homeostasis, we performed qPCR for a subset of inflammatory cytokines; unexpectedly we found there was a nonsignificant elevation of *Il22*, *Il23a*, and *Il6* at baseline in IEC *Casd1^-/-^*distal colons, suggesting a higher subclinical inflammatory tone (**Fig 2J**). To see if this was associated with any measurable impact on homeostasis, we analyzed baseline epithelial proliferation, which is sensitive to inflammatory stimuli^46^, by staining tissue sections with the proliferation marker Ki67 antigen; we observed no differences in proliferative status in distal colon epithelium (**Fig. 2K and L).** These findings suggest that under baseline conditions, epithelial *O*Ac-Sia deficiency may have a mild impact on basal inflammatory tone but does not affect the fundamental tissue organization and morphology of the intestine at any age.

### Impact of Sia *O-*acetylation on mucus function

Sialylation is necessary for complete mucus barrier function^16–18^, but the extent to which this depends on its *O-*acetylation is unclear. Indeed, current studies offer conflicting conclusions on the role of *O*Ac in Sia-dependent mucus function^37,40^. We therefore leveraged our IEC *Casd1^-/-^*model with complete *O*Ac ablation on mucus to address this question by conducting a detailed analysis of mucus structure on 3-month-old male and female mice through histochemical and lectin-based imaging and quantitation methods. By staining CFPE sections with Alcian Blue (AB), we found the fecal-adherent mucus barrier layer was similar when analyzed using cross sections of feces-containing distal colons of IEC *Casd1^-/-^* mice vs. cohoused sex-matched WT littermates (**Fig. 3A**). Because AB staining cannot distinguish between the proximally-derived niche and b1 layer, and distally derived b2 layer^10^, we examined if the barrier layer reflected any differences in these two key substructures in the context of *O*Ac loss. This was accomplished through dual staining with *Ulex europaeus agglutinin*-1 (UEA1; Fucα1,2-Galβ1,4GlcNAc-binding) and *Maackia amurensis* lecin II (MALII; 3*O*S-Gal/Siaα2,3Gal-binding), which showed the formation of the mucus barrier sublayers in both WT and IEC *Casd1^-/-^* animals. Analysis of mucus layer structure in colon tissue cross sections showed that in all investigated mice, clear and distinct b1 (UEA1^+^MALII^-^) and b2 (UEA1+MALII^+^) sublayers were formed (**Fig. 3B**). In addition, no difference in proximal colon mucus phenotypes was observed between the strains (**Fig. 3B**). These results suggest that Sia-*O*Ac deficiency does not affect fecal mucus encapsulation or the formation of mucus sublayers in the colon (**Fig. 3A and B**). We confirmed the intact barrier function of mucus by confocal imaging of mucus-microbe interactions using a universal bacterial probe EUB338 for fluorescence *in situ* hybridization (FISH) and mucus staining with the lectin UEA-1^1^, showing little microbial invasion into the mucus in the IEC *Casd1^-/-^* mice (**Fig. 3C**), which was seen in either sex. These studies explain why loss of *Casd1* does not lead to overt spontaneous inflammation at baseline.

**Figure 3.**
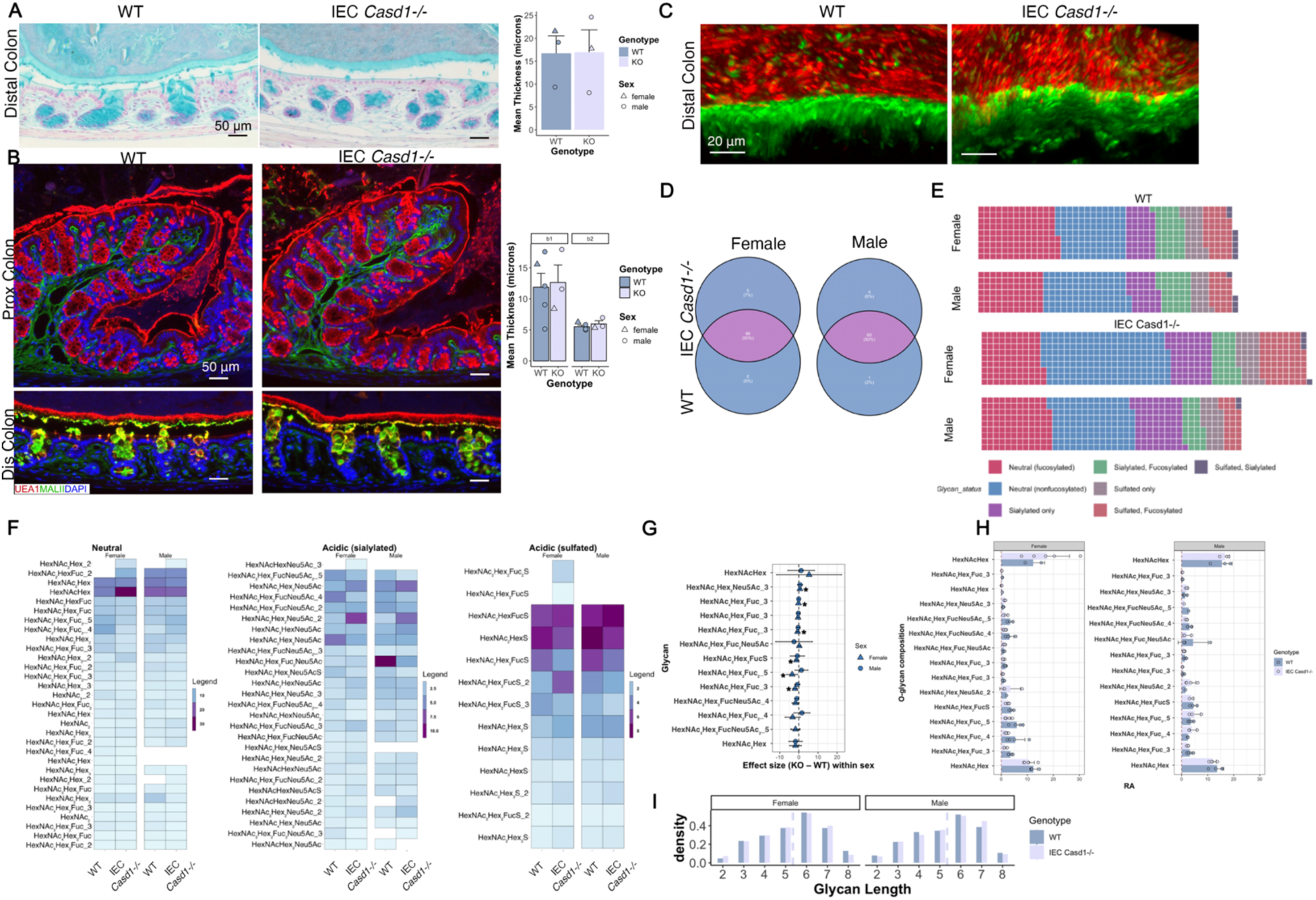
Loss of Sia-O-acetylation status does not impair mucus function. **A**. Representative Alcian Blue (AB) staining of CFPE distal colon sections showing overall *in situ* fecal-adherent mucus barrier structure. Right, bar plot (mean +/- SEM) overlaid with individual data points of mean mucus thickness per mouse. **B**. Epifluorescent imaging of lectin-stained (MALII, green; UEA1, red) CFPE sections of tissue and secreted mucus. Right, bar plot (mean +/- SEM) overlaid with individual data points of mean mucus thickness per male or female mouse as indicated. **C**. Representative (n = 4/genotype) confocal image of lectin-FISH-stained tissues from CFPE distal colon sections. **D**. Venn diagrams comparing similar (overlapping) and unique (non-overlapping) glycans between strains and sexes. **E**. Waffle chart showing relative abundances of individual classes of fecal Muc2-derived *O*-glycans between strains and sexes. **F**. Heatmap showing abundances of unique glycans in males and females within neutral or acidic groups. Absence of a tile indicates a non-detect in that specific group. **G**. Sex-stratified estimated marginal mean effect sizes (KO–WT ± 95% CI) for glycans selected by sex-adjusted main effects (*p*ₘₐᵢₙ < 0.05 or |estimateₘₐᵢₙ| > 1); asterisks indicate nominal significance (*p* < 0.05) from emmeans-based genotype contrasts within sex. **H**. Bar plot (mean +/- SEM) of relative abundances of specific glycans that were differentially abundant between WT and KO mice of either sex. **I**. Glycan length analysis depicted as histograms of frequencies of different glycan lengths, with the dashed line showing mean length.

### *O-*acetylation deficiency leads to subtle alteration in Muc2 *O*-glycosylation

Because loss of Sia *O-*acetylation exposes unmodified Sia to the lumen (**Fig. 2**), thus changing overall Muc2 glycosylation, we were interested to see how else the secreted mucin glycome was altered. We therefore analyzed the fecal mucus glycome from WT and IEC *Casd1^-/-^* mice using capillary LC/MS (in negative-ion mode)^1^ on mucus extracted from fecal pellets. Consistent with the stable mucus, we found the *O*-glycan profiles from IEC *Casd1^-/-^* mice showed no substantial differences from WT mice, though the relative abundance of some glycan classes slightly varied. 93% of glycan structures were common to both female WT and IEC *Casd1^-/-^* mice with 5% unique to IEC *Casd1^-/-^* and 0% unique to WT (**Fig. 3D**). Similar findings were observed in male mice, with 4% and 1% of glycans unique to IEC *Casd1^-/-^* and WT males respectively (**Fig. 3D**). To determine the influence of *O*Ac on relative abundances of specific glycan classes (Neutral, Sialylated, Sulfated, etc.), we parsed the glycome based on the sialylation, sulfation, and fucosylation status and compared the abundances using a waffle plot, which visually depicts proportions of detected *O*-glycans of a defined class. We found that neutral fucosylated glycans, as well as co-sialylated & sulfated glycans, were comparable among strains and sexes (**Fig. 3E**). However, male and female IEC *Casd1^-/-^* mice consistently had a greater abundance of “Sialylated only” glycans where Sia was the only capping structure (i.e., no other modifications such as fucose or sulfation), and female *Casd1^-/-^* mice uniquely showed increased abundance of extended glycans that were neutral with no capping fucose (“Neutral, non-fucosylated”), as well as co-modified sulfated + fucosylated moieties (“Sulfated, Fucosylated”) (**Fig. 3E**). Exploring at high resolution individual glycans revealed some differences. To simplify categories, we analyzed Neutral, Sialylated, and Sulfated structures, which revealed unique and common glycans found in male and female WT and IEC *Casd1^-/-^* mice (**Fig. 3F**). Interestingly, we observed some sex-specific differences in unique glycans, including a uniquely high abundance of a neutral HexNAcHex disaccharide in female IEC *Casd1^-/-^* mice, consistent with a core 1 structure (Galβ1,3GalNAcαSer/Thr) in female IEC *Casd1^-/-^* mice, which may reflect a partially degraded glycan. The effect of epithelial Casd1 loss on the *O*-glycome was assessed using estimated marginal means to account for sex-related variability and unequal representation of males and females across glycan species. Sex-stratified estimated marginal means revealed that differences in WT vs. IEC *Casd1^-/-^* vs across the glycome were modest, with substantial overlap between IEC *Casd1^-/-^* and WT mice and no glycans surviving FDR correction (**Fig. 3G**). However, a small number of glycans showed nominal (*p* < 0.05) genotype effects in females only, primarily involving fucosylated structures (HexNAc₂Hex₂FucS, HexNAc₂Hex₂Fuc_3, HexNAc₂Hex₂Fuc₂_5, HexNAc₄Hex₂Fuc_3, and the fucosylated isobar HexNAc₃Hex₂Fuc₂_3) which exhibited modest reductions in IEC *Casd1^-/-^* mice, as well as a single sialylated glycan, HexNAc₃Hex₂Neu5Ac_3, that was modestly increased (**Fig. 3G and H**). Notably, none of these effects were detected in males, and sex-stratified confidence intervals overlapped zero, indicating sex-specific variability rather than a consistent *Casd1*-dependent phenotype. Although the HexNAcHex (consistent with a truncated core 1 structure mentioned above) appeared to show a large effect suggestive of glycan degradation, the estimate was highly uncertain and not consistent between sexes, indicating that this difference likely reflects sampling imbalance rather than a true biological change. Consistent with this, a comparison of overall glycan length as a proxy for glycan degradation readout revealed no differences between WT and IEC *Casd1^-/-^* of either sex (**Fig. 3I**). Together, these analyses indicate that although loss of epithelial Casd1 profoundly reduces mucus sialic acid *O-*acetylation, it results in only subtle sex-dependent alterations in the broader mucin *O*-glycome, with no detectable impact on mucin barrier function.

### Role of Sia *O-*acetylation in regulating mucosal ecology

Given the complete absence of Sia-*O-*acetylation, and the role of *O-*acetylation in regulating sialic acid metabolism by bacteria^28,29^, we reasoned that loss of *O-*acetylation would drive changes in the microbiota community composition and function. We addressed this through next-generation sequencing of 16S rRNA amplified from fecal gDNA of WT and IEC *Casd1^-/-^*littermate mice, followed by both relative and quantitative microbial profiling (RMP and QMP, respectively) to gain a more comprehensive insight into the impact of *O-*acetylation on gut microbial ecology through measures of compositionality and microbial loads^47,48^. We first noted that loss of *O-*acetylation had minimal impact on β-diversity in male and female mice: Principal co-ordinate analysis based on Bray–Curtis dissimilarity revealed partial clustering but not by genotype or sex, indicating shifts in the relative abundance of dominant taxa by other factors (**Fig. 4A**). In contrast, weighted UniFrac did not show clear separation, suggesting that these differences occur primarily within closely related phylogenetic lineages rather than indicating broad phylogenetic restructuring (**Fig. 4A**). We also did not observe an impact on α-diversity as assessed by Shannon Index^49^ (**Fig. 4B**). We next analyzed total microbial load, estimated through qPCR (see Methods)^50–52^. Surprisingly, we saw a subtle but significant sex-specific impact of *O*Ac on total microbial loads, with only male IEC *Casd1^-/-^* mice showing a significantly reduced burden relative to male WT mice and female WT and IEC *Casd1^-/-^* mice (**Fig. 4C**). We next analyzed taxonomic differences through RMP and QMP. Analysis of the top 15 most abundant taxa in WT mice showed that genera *Duncaniella* (Phylum Bacteroidota), *Turicibacter* (Phylum Firmicutes), and/or *Prevotella* (Phylum Bacteroidota) contributed mostly to the overall relative abundance in both strains (**Fig. 4D**), although *Turicibacter* was consistently reduced in the IEC *Casd1^-/-^* samples vs. WT. As expected, the QMP profiles varied widely but were mostly within one order of magnitude of one another (**Fig. 4D**). We noted that particular species were proportionally different when comparing relative and absolute levels, including two genera of Phylum Bacteroidota: *Duncaniella* and *COE1*. We performed differential abundance analysis via MaAsLin2^53^ and identified seven taxa that met clear trend criteria (unadjusted *p* <0.05 and effect size >1) between WT and IEC *Casd1^-/-^* genotypes after adjusting for cage and sex: two *Turicibacter* species, a species of *Duncaniella*, and a species belonging to family Lachnospiraceae were enriched in WT mice, while two members of the *Kineothrix* genus and one member of the genus *Enterenecus* genus were enriched in IEC *Casd1^-/-^* mice. A more detailed statistical comparison of MaAsLin2-identified taxa to compare the distribution and means of their relative and absolute levels across genotypes was performed. Consistent with the idea that microbial load is a strong predictor of species abundance^48^, *Duncaniella* and *Turicibacter* dominated both RMP and QMP plots and both were reduced in IEC *Casd1^-/-^* vs. WT (**Fig. 4F**), in agreement with **Fig. 4E**. Species abundances varied over three orders of magnitude as shown by QMP data (**Fig. 4F**). Overall, the direction of changes in means of each taxon was similar between RMP and QMP (**Fig. 4F**); however, there was some discordance noted for one of the *Kineothrix* members in male mice (“g_Kineothrix_2”), one of the *Turicibacter* members (“g_Turicibacter”) and *Enterenecus* in female mice whose differences in relative abundances were not reflected in the absolute numbers, highlighting the utility of QMP to interpret RMP data^47,48^ (**Fig. 4F**). From the analysis of the MaAsLin2-identified taxa, it could not be determined whether these were responsible for the reduced overall load in the male IEC *Casd1^-/-^* mice relative to the other groups. Finally, we examined functional output through short-chain fatty acids (SCFAs). We found that loss of *O-*acetylation was associated with no major changes in SCFA levels, although sex-specific changes independent of genotype were observed, with male mice showing reduced overall SCFA production (**Fig. 4G**). Because *O-*acetylation is a natural inhibitor of neuraminidases^54^, we tested whether loss of epithelial Sia *O-*acetylation impacted fecal neuraminidase activity using the 4MU assay^23^. We observed no differences in neuraminidase levels (**Fig. 4H**), explaining why sialylated glycans were still abundant. In fact, determination of absolute levels of Sia on Muc2 glycans revealed a trend toward increased levels of Sia in IEC *Casd1^-/-^* males vs WT, although this was not significant (**Fig. 4I**). These studies suggest that while loss of epithelial Sia *O-*acetylation has a modest sex-specific impact on microbial loads, this does not translate to major changes in overall microbial community composition (with the exception of a few notable taxa) or SCFA production.

**Figure 4.**
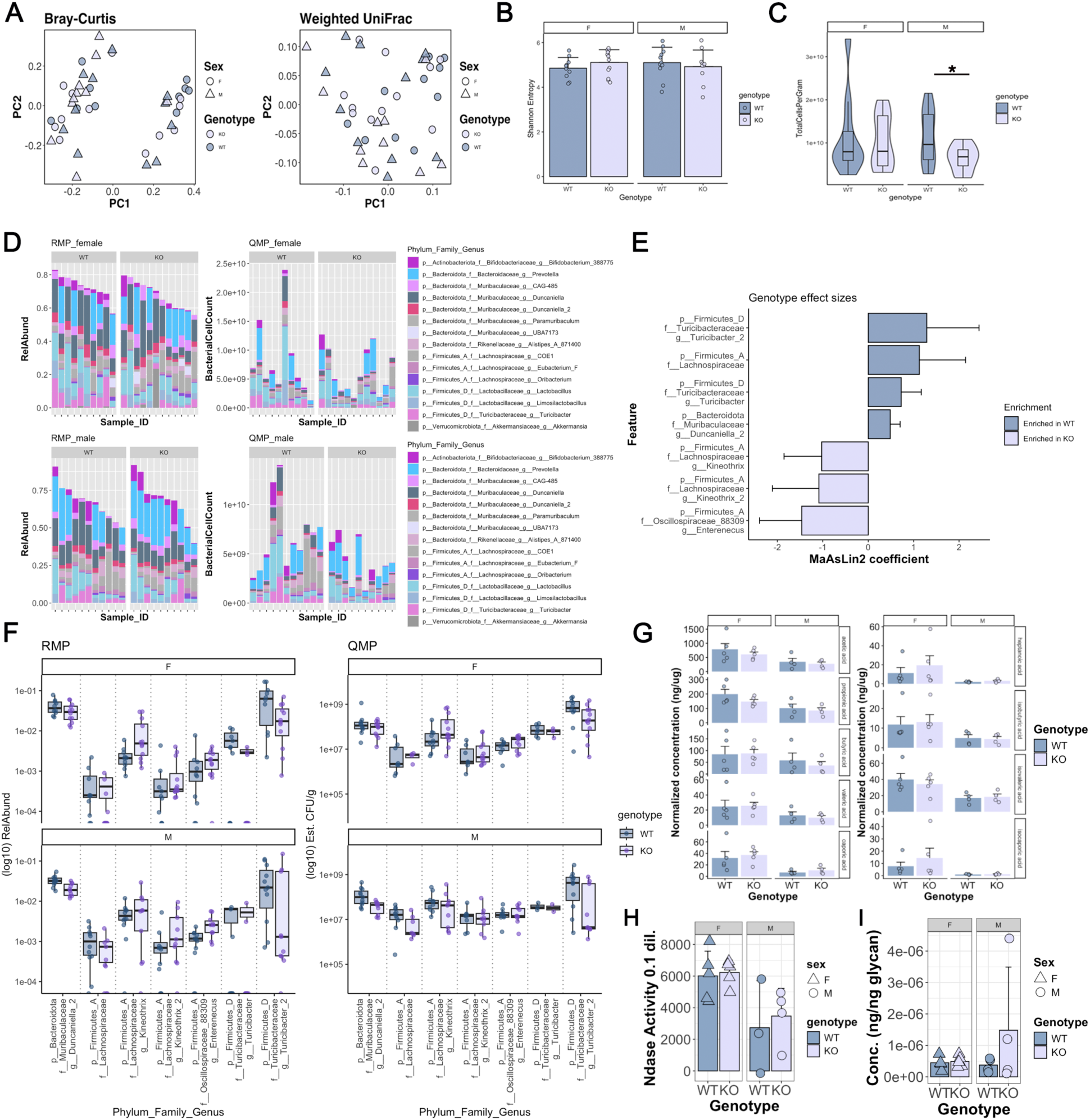
Impact of intestinal epithelial Sia *O-*acetylation on colonic microbial ecology. **A**. Principal coordinate analysis (PCoA) of Bray-Curtis dissimilarity and weighted UniFrac distances. **C**. Violin-boxplot showing estimated total bacterial loads per mg feces. Boxplot shows median + quartiles (\**p* < 0.05, Welch’s *t* test). **D**. Stacked bar chart showing the results of relative microbial profiling (RMP, right) and quantitative microbial profiling (QMP, left), of 15 of the most relatively abundant bacterial taxa in WT mice, and compared with the same taxa in IEC *Casd1*^-/-^ (KO) mice, with each lane representing an individual mouse. **E**. MaAslin2 analysis of differentially abundant taxa between WT and IEC *Casd1*^-/-^ mice determined from RMP values. **F**. Plots of relative (RMP) and absolute (QMP) values showing distribution of levels of MaAslin2-identified differentially abundant taxa (Genus level) comparing male and female WT and IEC *Casd1*^-/-^ (KO) mice. **G**. Bar chart showing normalized concentrations (mean +/- SD) of SCFAs per weight of WT or KO feces. Data points represent individual mice. **H**. Bar plot (mean +/- SD) units of fecal neuraminidase activities in feces. **I**. Bar plot (mean +/- SD) of Sia levels per ng of glycan pools.

### IEC *Casd1^-/-^* mice are more susceptible to acute intestinal injury vs. WT mice

Previous studies in mice lacking all epithelial sialylation due to conditional deletion of the CMP-Sia transporter Slc35a1 showed Sia was essential for protection from gastrointestinal insult from DSS^16^. To determine if epithelial *O-*acetylation was necessary for the ability of Sia to protect the mucosa, we performed a similar study with WT and IEC *Casd1^-/-^* mice. Mice were exposed to 2.5% DSS in drinking water for 7 days and analyzed for clinical and histological phenotypes (**Fig. 5A**). We found IEC *Casd1^-/-^* mice of both sexes showed a modestly more severe clinical response to 2.5% DSS vs. WT sex-matched mice, although with distinct dynamics: females showed increasing scores within the first 5 days, while male IEC *Casd1^-/-^* became worse after 4 days DSS (**Fig. 5B**). This could be attributed to a failure of female IEC *Casd1^-/-^* mice to gain weight over the first few days vs. WT females, and to a lesser extent, increased loss of body weight in male IEC *Casd1^-/-^* mice (**Fig. 5C**). At endpoints (day 7, DSS), we observed the large intestines of male IEC *Casd1^-/-^* exhibited moderately-increased damage from the insult vs. WT male mice (**Fig. 5D**). DSS-induced damage, including crypt dropout, inflammatory cell infiltrate, and ulceration, typically targets the distal colon in mice^55^; consistent with this, distal damage was observed in all mice, (**Fig. 5E–G),** but was greatest in male IEC *Casd1^-/-^* mice, with inflammation extending down into the submucosa in some animals (**Fig. 5E, G**). We also found injury extended into the mid and proximal regions of IEC *Casd1^-/-^* mice to a greater extent than in WT mice, in both sexes (**Fig. 5F, G**), which could further explain the trend toward higher clinical scores (**Fig. 5B**). Histologically, after adjustment for sex and colonic region, epithelial *Casd1* deficiency did not significantly affect the composite total pathology score (*p* = 0.17), although scores were consistently elevated in IEC *Casd1^-/-^* mice. However, epithelial Casd1 loss was associated with a significant increase in epithelial ulceration (β = +22.9, *p* = 0.029), with genotype-dependent exacerbation most pronounced in the mid colon (Δ = +37.3, *p* = 0.018) (**Fig. 5F, G**), indicating a colon-region-dependent protective capacity of *O-*acetylation against cytotoxin-induced epithelial damage. Mucus defects can exacerbate colitis^5,56^, and our past studies have shown that proximally-derived colon mucus forming the b1 barrier layer accounts for most of the spatial segregation of the colon microbiota via its primary encapsulating function around the fecal mass that it adheres to^10^, and that inflammatory damage in the proximal regions can disrupt this encapsulation and exacerbate colitis in more distally located colon regions in colitis-susceptible mice^10^. Therefore, we looked at the mucus layer in post-DSS-treated mice via Alcian Blue staining on CFPE sections in less-involved regions, showing more mucus disruption in the distal region of feces-containing IEC *Casd1^-/-^* mice. Using a linear regression model, we found the mucus layer was significantly reduced in thickness in IEC *Casd1^-/-^* of both sexes vs. WT mice (**Fig. 5H**) (*p* = 0.00138). Confocal imaging showed the mucus in IEC *Casd1^-/-^* mice was more degraded and had reduced barrier capacities toward the microbiota (**Fig. 5I & J**), explaining in part the tendency for worsened disease in the middle and distal colon in the absence of epithelial *Casd1*. Consistent with this, Spearman correlation analysis revealed distal mucus thickness was moderately inversely associated with both proximal colonic histologic damage scores (*p* = –0.39) and ulceration area (p = –0.47), indicating that thinner distal colon mucus layers were generally observed in mice with more severe proximal/mid tissue injury (**Fig. 5K**). The results suggest *O*-acetylated mucin from proximal regions, which is more diverse than in the distal colon (**Fig. 1**), serves to protect the proximal colon regions from cytotoxic injury, which in turn helps to preserve the middle- and distal colon-protecting b1 barrier layer. These studies identify intestinal epithelium as a major cell type through which *O-*acetylation protects the colon from insult.

**Figure 5.**
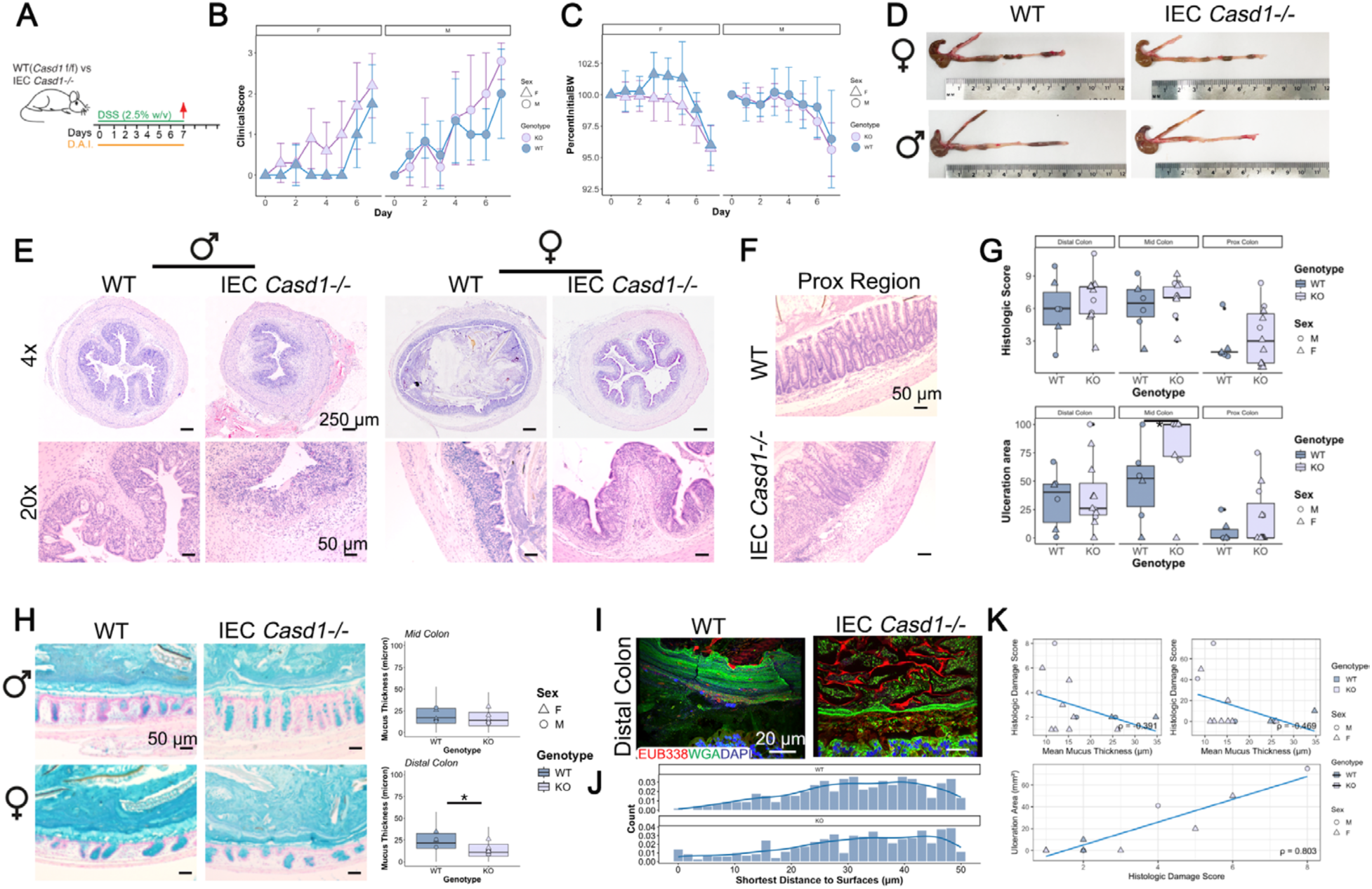
Loss of epithelial Sia *O-*acetylation exacerbates acute colitis caused by DSS. **A**. Schematic of DSS treatment strategy. **B**. Clinical disease activity after DSS treatment. **C**. Body weight loss after DSS treatment. **D**. Picture of colons of mice in response to DSS treatment. **E**. Representative low mag (upper panel) and high mag (lower panel) H&E staining of FFPE distal colons. **F**. Representative histology of FFPE colons at 7d post-DSS. **G**. Bar plot (mean ± SD) of histologic damage scores, with data points representing individual mice (\**p* < 0.05). **H**. AB staining of mucus in FFPE colons; Right, bar plot (mean ± SEM) of mean mucus thickness per condition, with data points representing mean mucus thickness of individual mice. **I**. Representative (n = 3 mice/group) confocal imaging of lectin:FISH-stained FFPE distal colons at 7d post-TM. **J**. Density plot quantifying numbers of FISH+ cells penetrating the mucus and tissues. **K**. Spearman’s correlation analysis between proximal colon damage and ulceration and distal colon mucus thickness (top plots respectively) and between ulceration levels and damage score as a positive expected correlation control (bottom plot).

### Intestinal epithelial Sia *O-*acetylation influences damage due to infection-induced colitis

We next sought to investigate whether *O-*acetylation played a role in the pathogenesis of a bacterium responsible for acute colitis. Previous studies have linked the bloom of pathogenic *E. coli* to the *O-*acetylation status of mucins *in vitro*^28^. *Citrobacter rodentium* is related to *E. coli* and belongs to a class of epithelial-adherent (non-invasive) pathogens called attaching/effacing (A/E) *E. coli*, similar to Enteropathogenic *E. coli* and Enterohemorrhagic *E. coli*. A/E pathogens cause infection by intimately adhering to the apical surface of epithelia in a type III secretion–dependent manner^57^. *C. rodentium* is also known to have intimate interactions with mucus^8,58^, and its virulence is impacted by Sia^22^. We therefore tested whether *O*Ac status impacted *C. rodentium* pathogenesis. We first inoculated WT and IEC *Casd1^-/-^* mice with *C. rodentium* and monitored both their stool counts and Clinical Activity Index over the next 10 days (**Fig. 6A**). Assessment of body weight and overall clinical disease activities (e.g., hunched posture, inactivity, diarrhea) did not exhibit any major differences between strains (or sexes); however, we did note a higher disease activity within the first 6 days in male IEC *Casd1^-/-^* mice (**Fig. 6B**), which correlated with reduced body weight vs. infected WT mice (**Fig. 6C**).

**Figure 6.**
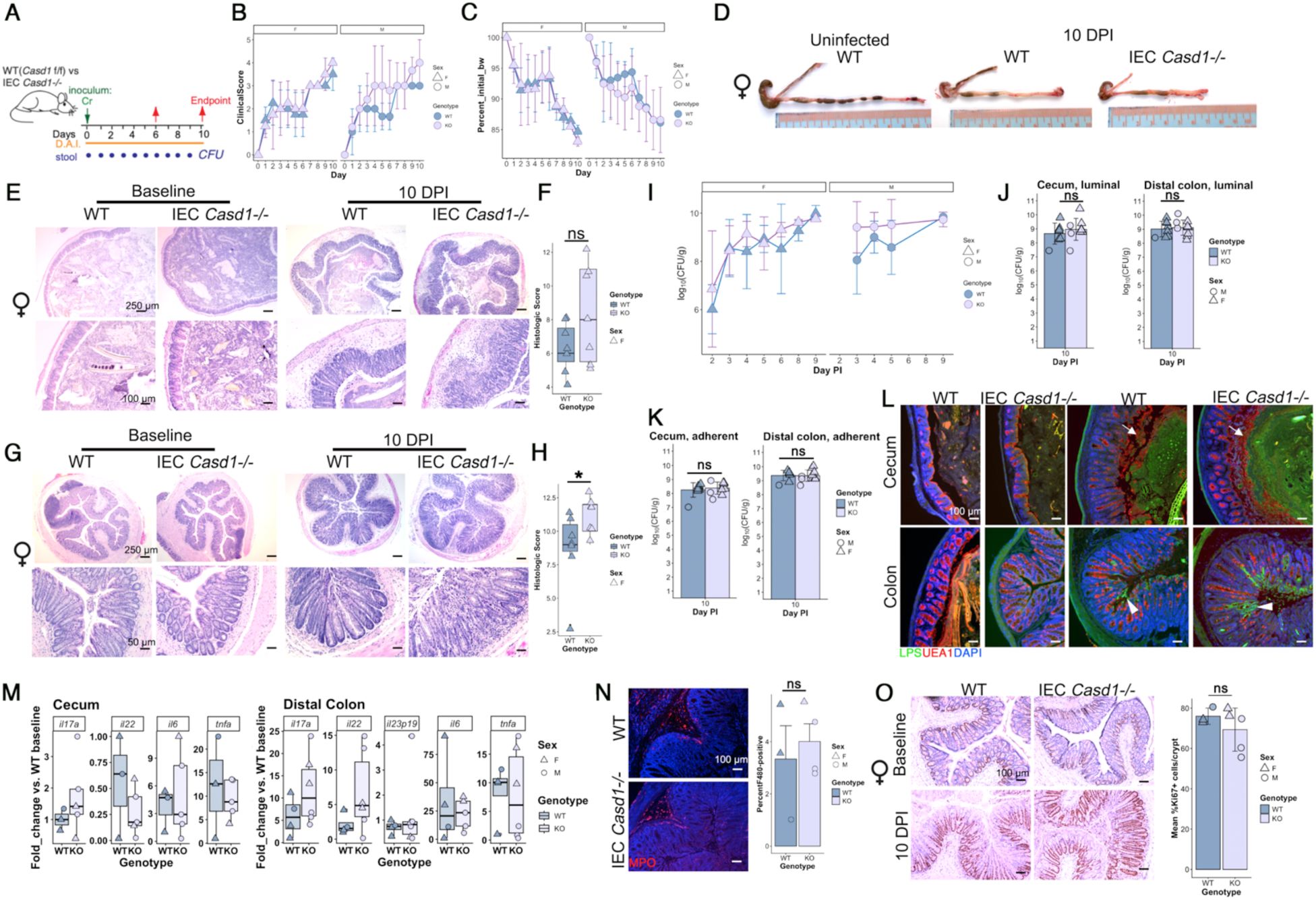
Epithelial Sia O-acetylation limits tissue damage but not colonization during *C. rodentium* infection. **A**. Schematic showing overview of the infection strategy. **B**. Clinical score stratified by sex. Results are plotted as mean ± SD. **C**. Longitudinal analysis of body weight following *C. rodentium* infection stratified by sex. Results are plotted as mean ± SD. **D**. Picture of female large intestine with terminal ileum. **E**. Representative low mag (upper panel) and high mag (lower panel) H&E staining of FFPE cecal tissues. **F**. Boxplot of histologic damage scores in cecum at 10 DPI, with points representing individual mice (ns, *p* = 0.228; rank-based ANCOVA). **G**. Representative low mag (upper panel) and high mag (lower panel) H&E staining of FFPE colon tissues. **H**. Boxplot of histologic damage scores in cecum at 10 DPI, with points representing individual mice (\**p* < 0.05; rank-based ANCOVA). **I**. Line plot of *C. rodentium* burdens (mean ± SD) in stool over time, stratified by sex. **J**. Bar plot (mean ± SD) of luminal *C. rodentium* burdens in the distal colon at experimental endpoint (10 DPI), with points representing individual mice. **K**. Bar plot (mean ± SD) of adherent *C. rodentium* burdens in the distal colon at experimental endpoint (10 DPI), with points representing individual mice. **L**. Epifluorescent labeling of LPS (green) and WGA (red) on FFPE tissues. Insets are magnified images of the corresponding boxed region. Arrows show secreted mucus. Arrowheads show mucosa-associated LPS typical of *C. rodentium* infection. **M**. Boxplot showing fold changes of inflammatory cytokine gene expression at 10 DPI relative to uninfected WT mice, as measured by RT-qPCR on RNA extracted from whole tissues. **N**. Representative IF staining for F4/80 on distal colon tissues. Arrows show influx of F4/80⁺ cells (macrophages) into the submucosa. Right: Bar plot (mean ± SD). Each point represents an individual mouse. **O**. Representative IHC for Ki67 on FFPE distal colon sections at baseline and experimental endpoint. *Right*: Ki67 index represented by a bar plot (mean ± SD) of fold changes in numbers of Ki67⁺ cells in infected vs. control crypts. Each point represents an individual mouse.

To examine how *C. rodentium* was influencing tissue damage, we first focused on 10 DPI, a time of fulminant infection with host responses underway^59^. Analysis of of large intestinal morphology at 10 DPI showed no obvious changes in the ileum in either group (as expected); however, infection led to shrunken and inflamed ceca that were more pronounced in male (**Fig. S2A**) and female IEC *Casd1^-/-^* mice (**Fig. 6D**), with female IEC *Casd1^-/-^* mice also showing increased thickness in the distal colon vs. female WT littermates (**Fig. 6D**). Histological analysis revealed infection led to heightened inflammatory damage to the cecum of infected mice of both strains; however, this phenotype was heightened in female IEC *Casd1^-/-^* mice at 10 DPI, which showed worsened submucosal edema and inflammatory cell infiltrate into the submucosa and mucosa vs. WT littermates at 10 DPI (**Fig. 6E, F**). Similar phenotypes were observed in the distal colon in female mice (**Fig. 6G, H**). Analysis of male mice at 10 DPI showed infection-induced cecal and colonic inflammation in WT and IEC *Casd1^-/-^* mice; however, differences in pathology between WT and IEC *Casd1^-/-^* littermates were not as stark as in female mice at this time point (**Fig. S2B, C**).

To gain insight into how epithelial Sia *O-*acetylation via Casd1 was influencing tissue pathology, we examined the relationship between *C. rodentium* colonization and disease activity, as these features are usually positively correlated^60^. Analysis of stool counts every other day following infection showed no significant difference in *C. rodentium* burdens in infected male and female IEC *Casd1^-/-^* mice (**Fig. 6I**). At 10 DPI, we analyzed luminal and adherent loads of *C. rodentium* in both the cecum and distal colon and found no overt differences in overall loads in either spatial compartment between male or female WT and IEC *Casd1^-/-^* mice (**Fig. 6J and K**). We also did not observe any major differences in systemic burden of *C. rodentium* between WT and IEC *Casd1^-/-^* mice of either sex, as assessed by colony counts in the spleen and liver (**Fig. S2D**). To gain spatial understanding of the relationship between disease pathogenesis and *C. rodentium* burdens, we performed immunostaining for *E. coli* LPS, which cross-reacts with *C. rodentium* LPS, along with the mucus marker wheat germ agglutinin (WGA), allowing us to determine spatial colonization patterns in relation to mucus between the strains. We found similar colonization patterns between WT and IEC *Casd1^-/-^* mice of either sex in the colon: LPS could be seen intimately associated with the apical surface epithelium in both strains in the distal colon only in infected mice (**Fig. 6L**; **Fig. S2E**). In the cecum, *C. rodentium* infection was associated with heightened secretion of mucin and more variable LPS signal on the epithelium, although it was more robust in the lumen (**Fig. 6L**; **Fig. S2E**). This suggested increased mucus defense in the cecum, which was confirmed by Alcian Blue staining (**Fig. S2F**), although no difference was apparent between WT and IEC *Casd1^-/-^* mice of either sex. Interestingly, in the cecum of WT mice at 10 DPI, increased 9-O-acetylated mucus was observed, as assessed by staining with the 9OAc probe (**Fig. S2G**), suggesting induced *O-*acetylation is linked to a tissue-protective response.

We performed a similar comprehensive analysis at 6 DPI. At this earlier timepoint, we observed significantly exacerbated disease in the cecum of infected male IEC *Casd1^-/-^* mice vs. WT littermates, and an obvious trend approaching significance in female IEC *Casd1^-/-^* mice vs. WT (**Fig. S3A and B**). As at 10 DPI, the histologic phenotype at 6 DPI was independent of *C. rodentium* burdens in the tissue or luminal compartments (**Fig. S3C and D**). *O-*acetylation status did not impact distal colonic disease phenotypes or burdens in male or female IEC *Casd1^-/-^* mice vs. WT mice at 6 DPI (**Fig. S3E–I**). These findings suggest that the impact of *O-*acetylation on susceptibility to pathogen-induced tissue damage is not due to regulation of *C. rodentium* numbers but rather host responses.

To gain insight into the underlying mediators of inflammation, we performed RT-qPCR for a panel of inflammatory genes known to modulate *C. rodentium*–induced pathology. In the colon, disease in IEC *Casd1^-/-^* mice was associated with elevated expression of *Il17a* and *Il22*, but not *Tnf*, *Il6*, or *Il23p19* (**Fig. 6M**). In the cecum, aside from moderately increased *Il17a*, there was no clear relationship between these cytokines and cecal disease (**Fig. 6M**), suggesting alternative mechanisms. Immunostaining for macrophage infiltration in the colon showed moderately higher numbers of macrophages in the submucosa of IEC *Casd1^-/-^* mice (**Fig. 6N**). The Ki67 index was similar between strains (**Figs. 6O and S2H**). Collectively, these studies suggest that intestinal epithelial *O-*acetylation plays a role in host defense by promoting tolerance to infection via limiting pathogen-induced damage rather than pathogen burden, although the mechanisms are likely complex and differ by tissue site.

## DISCUSSION

### Summary of novel findings of study

Our findings describe the first report of the role of epithelial sialic *O-*acetylation driven by Casd1 in gut homeostasis and protection from intestinal injury and insult. Although global *Casd1^-/-^* mice have been studied in the context of the gut^35^, the ubiquitous presence of *O*-acetylated Sia variants on glycoconjugates throughout the mammalian body limits our interpretation of how O-acetylated Sia impacts specific tissues, and what cells drive this response, in the context of a global deficiency. Our use of the IEC *Casd1^-/-^* mice overcomes these limitations, by controlling for potential stromal contributions of *O*Ac-Sia (e.g via Siglec interactions between immune cells^61^), and by defining the roles of key cell types through which Casd1 functions in the gut. We found that the detectable mucus Sia *O*-acetylome and its diversity are completely controlled by epithelial Casd1 in mice, but are not required for baseline homeostasis, or overall mucus function. However, *O-*acetylation does have important roles in regulating microbial ecology and protecting from acute intestinal injury caused by epithelial toxins and a mouse-adapted pathogenic bacterium, *C. rodentium*. These provide insight into why Sia diversity exists in intestinal epithelium at the level of *O-*acetylation and the impact its loss may have on colons of humans.

### Spatial distribution of OAc

*O*-acetylated Sia have been investigated on tissue scrapings or whole colon homogenates, but the spatial organization of *O*Ac on human mucus has remained unknown. Consistent with published reports^39,62^, through *in situ* studies with highly specific probes targeting *O*Ac-Sias and deep sialylomics on *in vivo* MUC2 extracted from healthy fecal material, we show that MUC2 is heavily *O*-acetylated and that this modification interacts with the endogenous microbiota. To our knowledge, this is the first investigation of sialic acid diversity on secreted human MUC2 *in situ*. While 9- and 7,9-*O*Ac were expected based on previous studies, we unexpectedly observed a 4-*O*-Ac signal at the edge of the human fecal mass. This modification is typically observed in equine, guinea pig, and fish samples^63–65^ and is not thought to be widely expressed on human mucins^39^. Consistent with this, sialylomics did not readily detect this variant. However, because it is difficult to resolve 4-*O*Ac by sialylomic approaches, we cannot rule out that 4-*O*Ac-Sia could be present on human MUC2. Based on its spatial relationship (above the mucus but within the mucus-associated microbiota), we are not certain this is part of the mucus and it may be associated with 4-*O*Ac-Sia–presenting microbes, possibly mucinophiles based on proximity to the mucus, although examples of this modification in the microbiota seem sparse. The presence of *O*Ac-Sia was conserved in mouse mucus, although proportionally less *O*Ac overall was observed compared to human MUC2. There was regional variation of this modification, with a notable absence in the cecum, both in secreted mucus and in goblet cells. The reason for this lack of Sia *O*Ac in the cecum is unclear. We also found that, to a small extent, the diversity and abundance of *O*Ac were inversely correlated with *Casd1* expression at proximal and distal sites. These studies point to region-intrinsic regulation of *Casd1* expression and activity that requires further investigation.

### Overall regulation of the O-acetylome and O-glycome

Our study definitively shows that while Casd1 does not modify the overall O-glycome to any major extent, *Casd1* is solely responsible for generating the *O*-acetylome of mucus in mice. Interestingly, in humans, the recently characterized *NXPE1*^36,37^, along with *CASD1*, contributes to Sia-9-*O-*acetylation. However, in mice, *Nxpe1* exists as a pseudogene^66^, explaining why no compensation by other potential *O*-acetyltransferases was occurring. The relative contribution of *NXPE1* and *CASD1* to the *O*-acetylome on mucus glycans in humans is unclear. Interestingly, in a recent genetic association study, there was a perfect positive correlation between mild peroxidase periodic acid–Schiff stain (mPAS)—a technique to distinguish non-*O*-acetylated (mPAS-positive vs. 9-*O*-acetylated (mPAS-negative)—and *NXPE1* expression but not *CASD1* expression histologically, suggesting *CASD1* is not sufficient in some human tissues to *O*-acetylate Sia in colon epithelium. However, we found that using the same assay, mPAS activities are entirely contingent on *Casd1* expression in mice (Fig. S2F). These data suggest there are possible regulatory differences of *Casd1* activities in human vs. mouse tissue. Biochemical assays of *NXPE1* also identified the generation of 9-*O*-acetylated variants of Sia *in vitro*. Whether *NXPE1* activity can lead to di- or tri-*O*-Ac variants (e.g., Neu5,7,9Ac; Neu5,7,8,9Ac) is not clear. *Casd1* appears to have the ability to *O*-acetylate C7 directly as well as C9. There is also evidence that supports migration of *O*Ac groups spontaneously to different carbons^31,67^, but whether this is happening with *NXPE1*-dependent *O-*acetylation is not defined. Furthermore, the contribution of *NXPE1* and *CASD1* to *O-*acetylation on human MUC2 *in vivo* is still unclear. It is likely both do, given they are expressed in the goblet cell Golgi apparatus. However, current evidence has looked either in tissues or in CRISPR-dependent *NXPE1*-depleted colon organoid systems, with the latter not directly showing the mucus structure^37^. With the new understanding that mucus can be readily analyzed non-invasively on feces, one could potentially resolve the contributions by identifying patients who have the *NXPE* G355R mutation^37^, collecting their feces, and analyzing *O*Ac status histochemically (e.g., mPAS) and/or via sialylomics. This would be important for future work.

### Overall impact of OAc status on baseline protection

The impact of *O*Ac-Sia loss on protection from insults were significant but unexpectedly subtle. The extent and diversity of *O*Ac on Sia would suggest an important physiological function *in vivo*. We obtained complete ablation of *O*Ac in gut epithelium and mucus in the IEC *Casd1^-/-^* model, which would be expected to leave Sia more susceptible to neuraminidase activities^25,28^. However, we observed similar amounts of Sia on mucus of the IEC *Casd1^-/-^* mice by glycomics and sialyomics, suggesting neuraminidase inhibition by *O*Ac is not a major regulator of Sia liberation in the colon. Although we did not measure Sia levels in the lumen directly, our results are consistent with the idea that while *O*Ac-esterases are already abundant in the colon ^68^, this does not appreciably render bound Sia more resistant to neuraminidases at baseline, whether derived from the host (NEU family ^40,69^) or microbe. However, caution must be made in this interpretation since we analyzed the barrier layer of mucus which does not house extensive microbiota; we currently cannot separately analyze the heavily colonized niche layer of mucus to compare the sialic content and see if there is a difference. Still, the observation that mucus is not extensively degraded in IEC *Casd1^-/-^* argues for a minor role of *O*Ac in regulating Sia levels on mucus glycans or on overall mucus integrity.

### Implications for disease

As observed for many pathologies, glycosylation of mucins and other glycoconjugates changes during disease states^70,71^. *O-*acetylation has been reported to decrease in inflammatory bowel disease^72^. This may explain why the cancer-associated glycan sialyl-Tn antigen is more readily observed via antibody detection in Ulcerative Colitis samples ^73,74^, since *O*Ac masks antibody binding. Thus, the reduction in *O*Ac-Sia may be secondary to the disease itself and not contributing directly to its etiology. Importantly there is an ostensible discrepancy over the biological roles of *O*Ac groups on Neu5Ac. In the study by Xavier and colleagues, the loss-of-function mutation is thought to be protective due to enhancing the barrier properties of the mucus, although this was not shown directly. In another study, inflammation-induced sialate *O*Ac esterase-dependent de-*O-*acetylation of Sia is thought to destabilize the mucus barrier and contribute to inflammation and inflammation-associated carcinogenesis^40^. The interpretation of both studies must be done with caution since the impact on mucus function was not directly addressed. What our studies suggest, at least in mice, is that *O*Ac loss has little impact on mucus integrity and homeostasis at baseline. However, our studies show that *O*Ac loss promotes more severe injury in response to acute toxin that is used as a model for IBD as well as to intestinal infection. These phenotypes suggest that at least in these contexts, *O-*acetylation of Sia is protective. It would be informative to combine the IEC *Casd1^-/-^* mice with models of spontaneous disease (e.g. IL10 KO) to determine if *O*Ac-Sia contributes to or protects from idiopathic inflammatory disease or cancer.

### Impact of OAc-Sia on microbial ecology

Since O-acetyl groups can inhibit microbial sialidases that would otherwise be liberated and accessed by Sia-utilizing microbiota, we expected a change in microbial ecology upon *O-*acetylation loss due to the potential for dysregulation of Sia metabolism; however, the change was not as dramatic as anticipated. No major changes in species composition or richness by beta- or alpha-diversity measures were observed. Still, some unexpected changes were noted, including a 2-fold reduction in total microbial load in male IEC *Casd1^-/-^* mice for reasons that are not clear. While calculating absolute microbial loads is still a relatively recent development in microbiome analysis, it is increasingly appreciated as an important variable when rendering conclusions on the impact of any intervention on microbiota dynamics^47,48,75^. Among the notable trends were those in the most abundant taxa, including *Duncaniella* (*family Muricibaculaceae, formerly S4-27*) and *Turicibacter*, both implicated in regulating gut health. For example, *Duncaniella* has two major relevant species. *Duncaniella muricolitica* has been implicated as a key modulator of the susceptibility of mice to colitis^76^. *D. muris* in contrast, is likely to be the isolate shown to ameliorate DSS colitis^77^. We are unclear which species are present in our facility. The relevance of *Turicibacter* is discussed below in the context of infection. Even with the shifts seen, they were not associated with any major functional changes as assessed by SCFA analyses between WT and IEC *Casd1^-/-^* and mice of either sex, or changes in neuraminidase activities. These studies indicate the impact of OAc-Sia on the microbiome is minor and not-likely to lead to any significant metabolic imbalances at baseline.

### DSS colitis

In the course of finalizing this manuscript, it was reported that global loss of *Casd1* led to similar increased susceptibility to DSS colitis, although the cell types and mechanisms have not yet been elucidated (as mentioned above)^35^. Our studies using the IEC *Casd1^-/-^* mice, where *O-*acetylation is preserved throughout the mouse except within the intestinal epithelium, strongly suggest that *Casd1* in epithelial cells represents the major cell type in protection from this insult. Our original goal in using this model was to address whether *O-*acetylation in the epithelium was the primary means by which epithelial Sia impacted susceptibility to disease ^16^. Our results argue this is not the case, due to the lack of spontaneous disease and relatively mild DSS-induced disease compared to the absence of epithelial Sia itself^16–18^. Notably, our studies suggest that *O-*acetylation of Sia has a regional influence, namely in the proximal and mid colon, which were more susceptible to injury by DSS compared to the same region in WT littermates. While the mechanisms underlying how proximal colonic *O-*acetylation is protective remain to be elucidated, our studies provide some key insights centering on colon mucus. First, proximal *O-*acetylation was far more diverse and abundant in the colon mucus compared to the distal colon (**Fig. 1**), suggesting a unique function of this modification within these regions. This is important since the proximal colon mucus at baseline is what forms the niche and barrier layers that directly interact with the microbiota^10^. Second, previous work has demonstrated that damage in the proximal colon correlates with defects in proximal colon mucus production that can impact its protective capacities in the distal colon^10^. Consistent with this, we found the mucus in IEC *Casd1^-/-^* mice during DSS was more extensively disrupted, with reduced thickness and barrier function in the distal colon. These studies are in line with mucus playing a role in reducing the severity of DSS colitis^5^. Thus, the ability of *O-*acetylation to protect the proximal colon region from damage by DSS allows the proximal colon region to keep producing barrier mucus that can further protect the distal colon from severe damage caused by cytotoxin and potentially infiltrating microbiota. Future research will need to elucidate the mechanistic relationship between the diversity of *O-*acetylation and its region-specific protective capacities, and whether something similar is occurring in humans.

### Implications for host defense

Overall, our results with *C. rodentium* indicated that epithelial *O-*acetylation was an important factor in limiting tissue damage caused by a biological insult. While both WT and IEC *Casd1^-/-^* mice developed disease from *C. rodentium*, it was consistently worse by histology in IEC *Casd1^-/-^* mice depending on sex and timepoint of infection. Curiously, male mice had worse cecal damage early on (6 DPI), but this normalized by D10 between strains. In contrast, female IEC *Casd1^-/-^* mice had consistently worse disease in the cecum and especially distal colon vs WT. In all cases, colonization was similar between the strains, so the increased damage cannot be attributed to microbial load differences. These findings implicate an increasingly appreciated type of host defense—namely “tolerance” defense, a harm-reduction strategy that limits tissue damage caused by the pathogen^78,79^. This is in stark contrast to more classic “resistance” defense strategies characterized by the deployment of direct microbial killing strategies like defensin production, reactive oxygen species, and phagocytosis^80^. This is in line with other functions of mucin-related glycans that have been reported to limit virulence of mucosal pathogens ^81–83^, or microbial communities ^84,85^. It is notable that while this was observed at the histological level, we did not note a strong correlation with clinical disease, suggesting the tolerance effect was mild. Nevertheless, several questions emanate from this work: For example, what is the mechanism of the increased damage in the IEC *Casd1^-/-^* mice? The susceptibility is consistent with recent studies showing *Turicibacter* spp abundance as negatively correlated with *C. rodentium*-induced inflammation^86^, as *Turicibacter* loads were overall reduced in IEC *Casd1^-/-^*mice. There may be an immunological component mediated by *O-*acetylation–dependent regulation of Siglec binding; however, this would predict the opposite of what we observe. In mice, Siglecs are predominantly inhibitory, and their binding is blocked by O-Ac Sia^25^. Thus, loss of epithelial Casd1 would be expected to enhance inhibitory Siglec engagement and dampen inflammation, rather than promote it. Alternatively, the free unbound Sia may also modulate *C. rodentium* virulence, since it is known that Sia is responsible for enhancing virulence gene expression^22^. However, the colonization patterns were not much different, suggesting this is not the major mechanism, or that more subtle changes are happening at the host-epithelial interface. It is notable that *C. rodentium* does not possess a neuraminidase^87^, so its ability to use liberated Sia would depend on either host or other microbial neuraminidase-dependent liberation. It would be interesting to test pathogens harboring neuraminidase activities, e.g., *Salmonella* Typhimurium^88^ or viral pathogens.

### Model Generation

As mentioned, conditional gene targeted systems are necessary to define the cells driving the function of widely expressed gene like Casd1. However, we encountered some unexpected challenges in an otherwise routine conditional model; rearrangement around the floxed allele. Cre-Lox technology has transformed functional genetic studies by enabling precise gene inactivation via site-specific recombination at loxP sites. However, floxed allele integrity is rarely assessed across generations, despite structural complexity that can lead to unintended genomic alterations. While on-target structural variants are well documented in CRISPR/Cas9 systems^89^, our findings suggest that similar rearrangements can also occur in Cre-Lox–based in vivo models. One plausible explanation is the occurrence of multiple *cis*-acting structural variants within the same engineered chromosome that can potentially arise from a concatemeric integration of the targeting construct at the *Casd1* locus. Tandem head-to-tail concatemeric insertions have been reported in mammalian genome engineering^90^ and shown to be prone to secondary rearrangements. In this context, increased structural complexity of the engineered *Casd1* locus may generate unstable recombination substrates which would explain WT-like allele restoration and emergence of additional floxed-derived amplicons. These studies raise important considerations on monitoring genomic integrity even when using established genetic systems.

### Conclusions

In summary, we report the first in vivo role of epithelial *O*Ac-Sia on mucus function and host defense in the intestinal tract. Our results indicate that while not necessary for intestinal immune homeostasis and mucus function, *O-*acetylation is a mechanism by which mucins fine-tune their interactions with gut microbiota and protect from inflammatory threats within the intestinal tract. Future studies will need to explore how *O*Ac-Sia impacts other chronic diseases of the gut including chronic inflammation and cancer.

## METHODS

### Animals

The “floxed” animals for the *Casd1* gene (B6(C3H)-*Casd1^tm1c(EUCOMM)Hmgu^*/PillJ, Stock No. 033573), along with male animals from a Cre transgene-expressing line (B6.Cg-Tg(*Vil1*-cre)997Gum/J, Stock No. 004586), were purchased from Jackson Laboratory. The Cre transgene in this line is expressed under the control of an epithelial-specific gene promoter, Villin, leading to Cre recombinase localization in the nucleus of intestinal epithelial cells (IECs). Genomic DNA extracted from ear notch samples obtained from these animals was subjected to genotyping to confirm the presence of the floxed and wild-type *Casd1* alleles, as well as the Cre transgene. Genotyping was conducted following JAX protocols (No. 36949 & No. 24364 for *Casd1* and generic Cre genes, respectively). Adult female *Casd1^f/f^* mice were bred with male Villin-Cre transgenic mice. Animals homozygous for the *Casd1* floxed allele (*f/f*) and carrying the Cre transgene (Cre⁺) were classified as conditional intestinal epithelial knockouts (IEC *Casd1^-/-^*mice). Other genotype combinations, such as *Casd1^f/w^*;VilCre⁺, *Casd1^f/w^*;VilCre⁻, *Casd^f/w^*;VilCre⁺/⁻, and *Casd1^f/f^*;VilCre⁻ were categorized as functionally wild-type (WT). In the majority of studies, we used *Casd1^f/f^*;VilCre⁻ mice as WT controls. The efficacy of deletion of the *Casd1* gene in epithelial tissues of male and female mice is described below. All animal procedures and experiments conducted in this study received approval from the Institutional Animal Care and Use Committee at the University of British Columbia (Protocol No. A19-0073) and adhered to the guidelines set forth by the Canadian Council on Animal Care. The animals were maintained on a standard diet (PicoLab Rodent Diet 20; LabDiet 5053).

### Deep Casd1 Genotyping

Three distinct primer pairs were designed using the Primer3 web service and validated to target specific regions within the retained sequence of *Casd1* transgenic cassette. The first primer set initially acquired from JAX designed to amplify a region encompassing the upstream LoxP sequence and certain portions of intron 1 of the gene (“Regular” Casd1 primers); the product amplicon of these primers excludes both exon 2 of the gene and the downstream LoxP sequence which are essential for knocking out the gene by Cre recombinase. The second primer set, referred to as E/D primers, was strategically chosen to amplify exon 2 of the *Casd1* gene (partial coverage) as well as the downstream LoxP sequence. The predicted PCR products were a 422 bp amplicon from the wild-type (WT) allele and a 452 bp amplicon generated from the floxed allele. A list of primer sequences is provided in **Table 1**. PCR amplifications were performed using genomic DNA extracted from ear notch samples. Subsequently, PCR products were separated by gel electrophoresis on a 3% agarose gel. Gel images were analyzed to confirm the presence of expected amplicon sizes corresponding to the intact transgene construct and the absence of unintended deletions or rearrangements. Individual PCR amplicons were extracted from the agarose gel using a QIAquick Gel Extraction Kit following the manufacturer’s instructions and subjected to Sanger Sequencing.

**Table 1.**
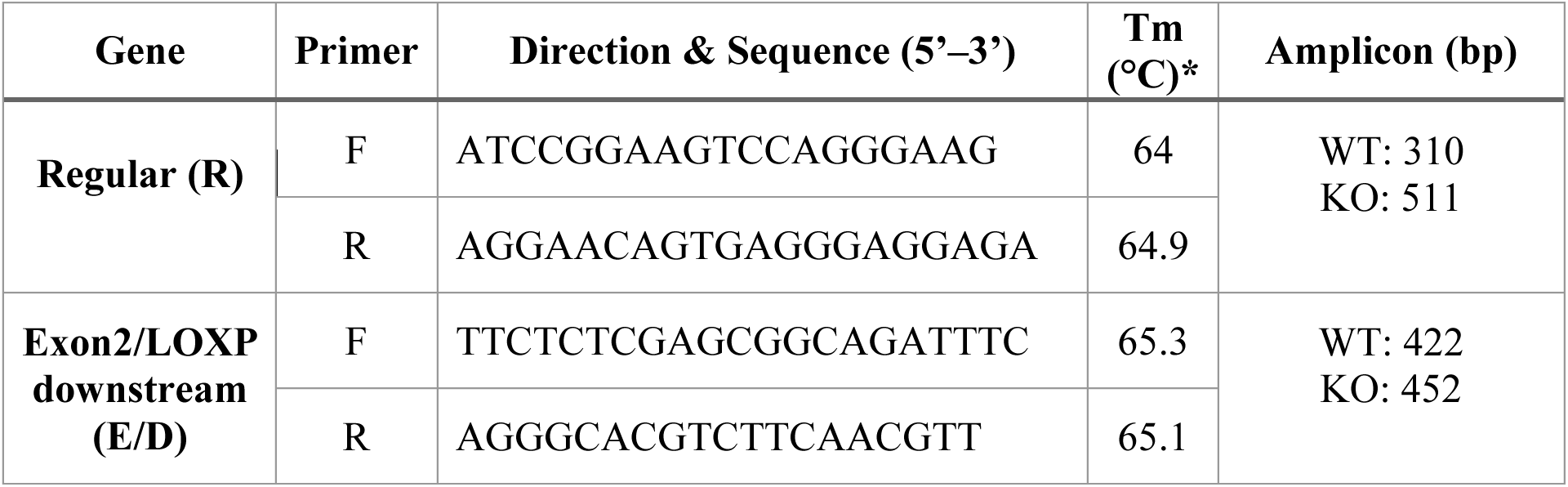
Primers used for deep genotyping.

### Dextran sulfate sodium (DSS) treatment

WT and IEC *Casd1^-/-^* (male and female) co-housed littermates were treated with 2.5% DSS (∼40 kDa; a 1:2 mixture from MMP Biologicals (USA) or TdB labs (Sweden)) dissolved in drinking water and filter sterilized (0.2 µm) for 7 d. Mice were then euthanized for downstream analysis of mucus quality and histology of the colon. Body weight, the presence of occult blood per rectum, and stool consistency were measured daily during the course of treatments to obtain a clinical score as previously described^15^. Histologic scoring is described below (“Histology”).

### Bacterial strains and infection of mice

Mice were infected by oral gavage with 0.1 ml of an overnight culture of LB containing approximately 2.5×10^8^ cfu of *wt C. rodentium* (formerly *C. freundii* biotype 4280, strain DBS100). For bacterial enumeration studies, a streptomycin-resistant derivative of *C. rodentium* DBS100 was utilized.

### Tissue Collection and Processing

Adult WT and IEC *Casd1^-/-^* littermate mice (8–16 weeks old) were euthanized, and their colons were extracted. Approximately 1 cm segments from the proximal and distal colon were cut using a single-edge disposable blade and placed in Carnoy’s fixation solution (60% methanol, 30% chloroform, 10% glacial acetic acid) for either a minimum of 2 h at room temperature or 12 h at 4 °C. Samples were then processed overnight before paraffin embedding using a Leica TP 1020 tissue processor. The following day, the samples were oriented vertically to obtain cross-sections and subsequently paraffin-embedded (Leica Histoscore Arcadia H). Carnoy’s-fixed, paraffin-embedded (CFPE) colon tissues were sectioned at 5 µm thickness (Micron HM 355S, Thermo Fisher), and every two sections were collected onto separate positively charged slides.

### Histochemical Staining and Quantitation

*Alcian Blue*. CFPE sections were cut (5 µm), deparaffinized and hydrated using standard protocols. Sections were then immersed with Alcian blue (AB) pH 2.5 (Sigma), which stains acidic mucins a light blue^91^, for 20 min and then thoroughly rinsed in tap water. Sections were counterstained with nuclear fast red for 10 min, dehydrated and mounted with Permount and imaged using a EVOS M5000 under brightfield settings.

#### Mucus thickness measurement

Mucus thicknesses were measured using the InterEdgeDistance Macro (V3) (https://forum.image.sc/t/imagej-macro-to-measure-distance-between-two-user-drawn-lines-edges-version-3/110803). For DSS treated sections, we noted the AB-stained mucus was disrupted, and therefore employed an in-house script to accommodate the variability and quantify the mucus layer. Mucus thickness was quantified using a custom ImageJ/Fiji macro developed for analysis of degraded mucus in DSS-treated mouse distal colon cross-sections imaged at 4X magnification. Images first underwent colour deconvolution using the Alcian Blue/Nuclear Fast Red vector to separate mucus and tissue signals, with the tissue channel used to detect the epithelial surface. The curved tissue–mucus interface was then identified and straightened using ImageJ’s Straighten function. Adaptive thresholding (Intermodes method) was applied to segment the faint barrier mucus layer while excluding the denser niche mucus. The upper and lower mucus boundaries were manually defined, and thickness was measured at 1-pixel intervals across the straightened image, converted to micrometer using a calibration of 0.6467 px/µm, and summarized as mean, standard deviation, minimum, maximum, and coverage percentage, and results were exported as CSV files for downstream analysis.

#### PB–KOH–PAS

*O-*acetylation of Sia at C7 and C8 was detected by the PB–KOH–PAS method ^45^. CFPE tissue sections were deparaffinized and dehydrated, then treated with 1% sodium borohydride in 1% disodium hydrogen phosphate for 30 min. After washing with water, tissues were incubated with 0.5% KOH in 70% ethanol for 30 min, followed by a 70% ethanol wash and a rinse in tap water. Sections were oxidized in 0.5% periodic acid for 5 min, rinsed in distilled water, and placed in Schiff reagent for 15 min. Sections were washed in tap water (5 min) and counterstained with Gill’s hematoxylin for 1 min. Tissue sections were mounted with Paramount mounting medium (Fisher Scientific). For PB-PAS negative controls, the KOH step was omitted.

#### Mild peroxidase PAS (mPAS)

For mPAS staining, the procedure was performed as described^36^. Briefly, deparaffinized, rehydrated sections were incubated in 0.1 M acetate buffer (pH 5.5; Thermo Fisher, AM9740) at 4°C. Oxidation was performed with 1 mM (0.02%) NaIO₄ (Sigma Aldrich, P7875-100G) in acetate buffer, followed by treatment with 1% aqueous glycerol (Sigma Aldrich, G5516). Slides were then treated with Schiff’s reagent (Sigma Aldrich, 3952016-500 ML), rinsed in 0.5% K₂S₂O₅ (Sigma Aldrich, 60508) in 0.05 M HCl, washed repeatedly in distilled and tap water, dehydrated, mounted, and imaged as above.

### *In situ* analysis of Sialic Acid *O-*acetylation

To study *in situ* 9-*O*Ac expression, sections were air-dried overnight, then dewaxed in xylene and rehydrated. The sections were permeabilized in (0.05% Tween 20 in PBS; 10 minutes at RT), followed by blocking with streptavidin and biotin solutions (Vector Laboratories) for 10 minutes each at RT, prior to immunostaining. The 9-*O*Ac–specific Porcine Torovirus P4 esterase (PToV HE), the 7,9-*O*Ac–specific Bovine Coronavirus Mebus strain esterase (BCoV-M HE), and the 4-*O*Ac–Sia–specific Mouse Hepatitis Virus S strain *O*-acetylesterase (MHV-S HE). These inactive esterases preserved binding but not removal of *O*Ac, thus permitting their use as probes. Probes with active esterases or had point mutations that inhibited binding (e.g. PToV-P4 HE-Fc, F271A for 9-*O*Ac) were used as negative controls to confirm specificity. Probes were diluted 1:100 (final concentration of 0.5 µg/mL) in blocking solution (0.1% BSA, 0.05% Tween 20 in PBS). Tissues were then incubated overnight at 4°C. The following day, the primary antibody was washed 4X with PBST. Tissues were then incubated with biotinylated goat-anti- human IgG antibody (Vector Laboratories) diluted to 2:250 (final concentration of 0.5 µg/mL) for 2 h at RT. Following three washes with PBST, slides were incubated with Alexa Fluor™ 350-, 488-, or 594-conjugated Strepatividin (Invitrogen) diluted to 1 ug/ml in ADB for 45 minutes at RT. Slides were washed in PBS and counterstained with DAPI (2 ng/mL in dH2O) water for 3 minutes at RT followed by four washes in water. Finally, slides were mounted with Fluoroshield mounting media (Sigma, SKU F6182-20ML). Tissues were imaged using a EVOS^TM^ M5000 microscope (Invitrogen) at 10X and 20X magnification.

### Immunostaining and Image Analysis

Formalin- or Carnoy’s-fixed paraffin-embedded sections were deparaffinized and rehydrated. For streptavidin–biotin-based staining, sections were blocked using Streptavidin/Biotin Blocking Solution (Vector Labs). For immunohistochemistry (IHC), endogenous peroxidase was quenched with 3% H₂O₂ for 10 min at RT. For epifluorescence-based immunolabeling or lectin-based staining, Serum-Free Protein Block (DAKO) was used (10 min, RT). For F4/80 labeleing, sections were incubated with AF594-conjugated rat anti-murine F4/80 (details) overnight, counterstained with DAPI, and mounted. For dual LPS–mucin labeling, sections were incubated with rabbit anti-*E. coli* LPS (2 µg/mL) for 2 h at RT, washed, then stained with rhodamine-conjugated WGA (2 µg/mL, Vector Laboratories) and detected with DyLight 488–conjugated donkey anti-rabbit IgG (5 µg/mL; Jackson ImmunoResearch) for 1 h at RT. Slides were counterstained with DAPI (20 ng/mL, 2 min, RT in the dark) and mounted. For lectin labelling**, s**ections were incubated with biotinylated *Maackia amurensis* lectin II (MAL-II, 1:200, Vector B-1265) for 2 h at RT or overnight at 4°C, rinsed with PBS, and incubated with DyLight 488–Streptavidin (5 µg/mL; Jackson ImmunoResearch) and rhodamine–UEA-I (2 µg/mL, Vector Laboratories) for 1 h at RT in the dark. Slides were washed three times in dH₂O, stained with DAPI (20 ng/mL, 5 min), washed, and mounted with Permafluor. For Ki67 staining, sections were incubated with polyclonal rabbit anti-Ki67 (Thermo Scientific), washed, incubated with HRP-conjugated goat anti-rabbit IgG (2 µg/mL; Jackson ImmunoResearch) for 40 min at RT, and developed using DAB (Vector). Sections were counterstained with hematoxylin, dehydrated, and mounted. Images were taken on an EVOS system as described above. Species-specific isotype controls or omission of primary antibody were used as negative controls.

#### B1/b2 mucus thickness measurement

Mucus thickness was quantified manually in ImageJ from fluorescence-stained tissue sections by measuring the perpendicular distance between defined fluorescent layers. Using 20X images, line measurements were drawn from the epithelial surface to the outer edge of the b2 (MALII+UEA1+) mucus layer and separately from the top of the b2 layer to the outer edge of the b1 (MALII^-^UEA1+) mucus layer to get b1 thickness, with 25 – 30 measurements taken per section to account for regional variability. Thickness values for each layer were recorded in pixels, and then converted to microns using a scale derived from a known length of a scale bar for that image. Raw data was imported into an excel spreadsheet and processed in R for downstream visualization and statistical analysis (described in Statistics)

#### Enumeration of proliferating cells

For proliferation analysis, 10–20 well-oriented crypts (visible crypt orifice and crypt base) were analyzed per section. In each crypt, the number of Ki67^+^ nuclei were enumerated per crypt. The average number of Ki67^+^ cells per crypt was determined for each mouse, and the mean of these averages was determined among all mice in each respective experimental group.

#### Enumeration of Macrophages

To enumerate infiltrating macrophages into tissues, we developed an ImageJ Macro that performs automated quantification of macrophages in immunostained tissue sections by identifying F4/80-positive cells and normalizing them to total cell number. In brief, the blue (DAPI) channel is processed to generate a binary nuclear mask using contrast enhancement, noise reduction, automatic thresholding, and watershed segmentation, followed by particle analysis to count total nuclei. In parallel, the red (F4/80) channel is thresholded and analyzed to quantify total F4/80-positive objects. Co-localization is then assessed by intersecting the F4/80 and DAPI masks, allowing enumeration of nucleated F4/80-positive cells as a proxy for bona fide macrophages. For each image, total nuclei, total F4/80 signal, nucleated F4/80-positive cells, and the ratio of macrophages to total nuclei are recorded in a CSV file, and labeled mask images for DAPI, F4/80, and their overlap are saved for quality control.

### Mucin-FISH labeling and quantitative imaging

Briefly, in order to perform the fluorescence *in situ* hybridization (FISH) staining process, fecal pellet sections were initially deparaffinized and rehydrated, then treated with FISH buffer (20 mM Tris-HCl, pH 7.4, 0.9 M NaCl, 0.1% SDS) for 5 minutes at room temperature (RT). The sections were then incubated with a Texas Red–conjugated universal bacterial probe EUB338 (5’-GCTGCCTCCCGTAGGAGT-3’; bp 337–354 within bacteria EU622773; Eurofins MWG Operon) and a genus-specific probe for *Prevotella* and *Bacteroides*, BAC303, diluted 1:100 in FISH buffer overnight at 37 °C^92^. After the incubation step, sections were washed with FISH buffer (1X, 5 minutes) and PBST (2X, each for 5 minutes), and then incubated with an LS anti-MUC2 antibody (Gift from Chadee) diluted 1:500 in antibody dilution buffer (AbDB) (0.1% BSA, 0.05% Tween 20 in PBS) and/or the *O*Ac-Sia-targeting probes described above for 2 hours at RT. This was followed by secondary staining using Donkey Anti-Rabbit IgG conjugated with Alexa Fluor® 594 (final concentration of 5 µg/ml in AbDB) to achieve fluorescent staining of the mucus layer. The sections were then counterstained with DAPI (1:2000 in H₂O) and mounted before imaging. In this approach, bacteria have been visualized within the gut lumen and their proximity and localization relative to the mucosal surface were measured and compared between WT and IEC *Casd1^-/-^* mice. The stained images were imaged either by eplifluorescent microscopy with the EVOS M5000, or by confocal microscopy using an Olympus FluoView FV1000 microscope. Z-stacked confocal images were acquired with a 20X dry objective, or a 60X oil-immersion or 40X dry objective, with optical sections taken at 0.3 to 0.5 μm intervals. Images were analyzed using Imaris software (versions 7.7.2 and 9.5) as described below.

#### Bacterial distance from mucosal surface or bottom of mucus

For confocal images stained with FISH probes and mucus markers (lectins or Muc2 antibody, described above), the distance of each bacterium (∼300–2000/image) to the mucosal surface (or bottom of mucus for fecal mucus studies) was determined using the Surfaces function within Imaris software. Briefly, Imaris Surfaces accurately (∼90–95% precision) pinpointed each bacterial cell and assigned them a surface and ID. A separate surface (called “Tissue”) was generated for the mucosal tissue, either automatically or via careful manual drawing of contours along a clearly visible mucosal surface. The software then calculated the distance of each bacterium to the mucosal surface using the “distance to nearest surface = Tissue” calculation. The calculations were verified using a subset of manually measured bacteria. Data were exported into Microsoft Excel and then to GraphPad Prism to generate a histogram of the frequency distribution of the absolute numbers of bacteria at a given distance from the mucosal surface, or in the case of fecal sections, the bottom of the mucus layer.

### Colon crypt isolation

Crypt isolation was performed according to established methods^15^. 1.5 cm-long pieces taken from longitudinally-opened and PBS-washed distal colons were cut into 3 – 4 pieces, and placed into holding solution (1% FBS, 1% Penicillin/Streptomycin, 50 mM HEPES pH 7.4 in 1x HBSS) on ice until all tissues were harvested. Tissues were simultaneously placed in tubes containing 1 mL crypt isolation buffer (2.5 mM EDTA, 2 mM DTT in HBSS), and incubated with moderate rocking for 40 minutes at 4°C. The tissue were allowed to settle, the supernatant was collected, and fresh isolation buffer was added and the isolation step was repeated 4 more times. After 4 mL was collected, the procedure was repeated 4 more times but with vortexing pulses (2 x 5s) in between washes, until a total of 8 mL was collected. Supernatants were then centrifuged at 150g for 5 minutes at 4°C. Supernatant was aspirated off, and pellet was washed in 200 – 400 μl of sterile cold PBS. Pellets were then re-centrifuged, supernatant removed, and stored in 80°C until used. Viability was 80 – 90% as determined by trypan blue. The remaining tissues were taken for protein and nucleic acid extraction as below.

### Extraction of protein and nucleic acids from IECs and tissues

Distal colon tissues ∼ 1 cm^2^ were placed in 1 mL of Trizol® with 2 mm metea beads (Qiagen) and homogenized for 1 min at 30 HZ using a Retch Bead miller. IECs were given 0.75 mL of TRIzol LS per 5 × 10⁶ IEC and did not undergo bead milling. Phase separation was performed by addition of 200uL of chloroform/0.75mL of TRIzol LS, after which the RNA-containing aqueous phase was processed through isopropanol precipitation, ethanol washing, and resuspension in storage buffer (RNAase-free water, 0.1mM EDTA, and 8mM NaOH or 1x TE) before storage at -80°C. DNA was isolated from the interphase and organic phase using ethanol precipitation and sodium citrate washes (10% EtOH and 0.1M sodium citrate), followed by ethanol dehydration and dissolution in storage buffer prior to -80°C storage. Protein was subsequently isolated from the remaining supernatant by isopropanol precipitation using 0.75mL of 100% isopropanol, followed by incubation, centrifugation, guanidine hydrochloride in ethanol washes (1.5mL of 95% Ethanol with 0.3M Guanidine hydrochloride), and final ethanol washing and drying. Protein pellets were solubilized in 8M Urea and 1% SDS with sonication before storage at -80°C. These purified fractions (RNA for RT-qPCR of the Casd1 gene, DNA for genotyping, and protein for 9-*O*Ac probing and western blot) were used to confirm *Casd1* knock out and loss of 9-*O*Ac expression in IECs extracted from colon tissues.

### RT qPCR

The concentration and purity of total RNA extracted from the isolated IECs or tissues were evaluated using a spectrophotometer. Subsequently, cDNA synthesis was conducted using a commercially available reverse transcription kit (Omniscript RT Kit, Cat. no. 205111). Briefly, RNA samples were treated with DNase to eliminate genomic DNA contamination, after which 1 µg of RNA was converted into cDNA using the reverse transcription kit and oligo(dT) and random hexamer primers, as per the manufacturer’s instructions. The cDNA product from the previous step was utilized as template for quantitative real-time PCR (RT-qPCR) using specific primers for the *Casd1* gene. PCR reactions were performed in duplicate, and a non-template control (NTC) PCR was included in each round of experiment to rule out contamination with genomic DNA.

Gene expression for various genes (**Table 2**) was measured using a CFX Opus 96 Real-Time PCR system (BioRad). PCR amplification was performed in a final volume of 10 μl, containing 1X DNA polymerase, 1.5 mM MgCl₂, 1 mM KCl, 250 ng of cDNA, 1X SYBR Green master mix, and 0.6 μM of each primer. qPCR was carried out according to the conditions in Table 1. The Cq (quantitative threshold value) was obtained, and exported into R and gene expression was calculated and plotted with an in-house R script using the ddCt method and comparing fold-change to WT controls at baseline. The data were reported as mean ± S.D.

**Table 2.**
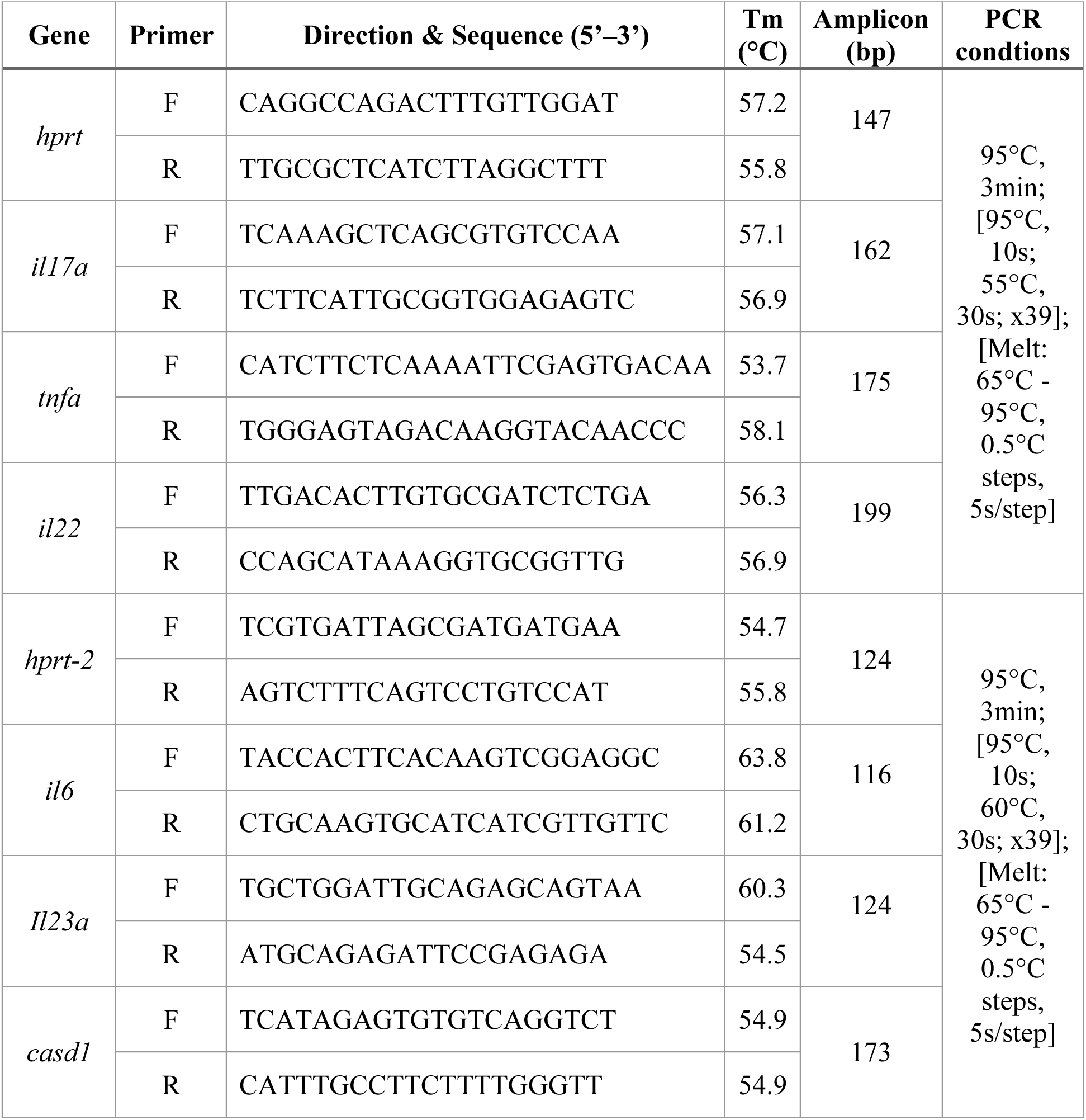
Primers and qPCR conditions used in this study.

### Gel electrophoresis and western blotting

Proteins extracted from crypt isolated cells were analyzed using composite sodium dodecyl sulfate urea agarose polyacrylamide gel electrophoresis (SDS-UAgPAGE). The composite gel comprised a gradient of 0.5 to 1.0% agarose, 0 to 10% glycerol, and 0 to 6% acrylamide. The running buffer used was buffered boric acid (192 mM boric acid, 1 mM EDTA, 0.1% SDS, pH 7.6). Protein samples were mixed 1:1 with 2x Laemmli Sample Buffer (2x SB), boiled for 5 minutes, and loaded onto the composite gel. Electrophoresis was performed for 2 hours at 110 V on ice. Post-electrophoresis, proteins were transferred onto a polyvinylidene difluoride (PVDF) membrane via wet transfer at 100 V for 1 h on ice using standard Tris-Glycine transfer buffer with 20% methanol. The PVDF membranes were briefly washed with TBST and then blocked with 2% (w/v) BSA in TBST (10 mM Tris, pH 8.0, 150 mM NaCl, 2% Tween-20) for 1 hour at RT. This was followed by blocking with a Streptavidin-Biotin blocking kit (Vector Laboratories) according to the manufacturer’s instructions. Membranes were incubated ON at 4°C with the 9-*O*Ac probe (PToV-P4 HE-Fc) at a final concentration of 0.1 µg/ml in TBST. After 6X 10-minute washes in TBST, membrane was incubated with biotinylated goat-anti-human IgG antibody (Vector Laboratories) at a final concentration of 0.25 µg/ml for 2 h at RT. Following another 6X 10-minute washes in TBST, membrane was incubated with HRP-conjugated streptavidin diluted 1:4000 in TBST for 1 h at RT. The membrane then underwent a final 6X 10-minute washes in TBST before immunoreactive bands were detected using Pierce SuperSignal Enhanced Chemiluminescence reagent (Thermo Scientific) on the ChemiDoc MP imaging system (BioRad).

### Clinical Scoring

At baseline, and during induced diseases states, mice were analyzed for clinical manifestations of disease, measuring parameters related to spontaneous colitis, including weight loss, stool consistency, rectal prolapse, bleeding, activity, and pain symptoms like piloerection. The grading system used was described in detail in previous publications^14^. Weekly assessments were performed, and mice were evaluated until 50 weeks of age.

### Histological Scoring

Histological scoring was performed according to established parameters for chemically induced colitis and bacterial-induced colitis described in Erben et al^93^. Briefly, distal colonic inflammation was evaluated on formalin-fixed, Gill-hematoxylin and eosin-stained distal and mid colon tissue sections using a semi-quantitative composite scoring system assessing inflammatory cell infiltration, mucosal architectural injury, mucosal bleeding, and lesion area. Inflammatory cell infiltration was graded as 0 (absent), 1 (mild, confined to the mucosa), 2 (moderate, involving mucosa and submucosa), or 3 (severe, transmural). Architectural injury was scored as 0 (normal), 1 (focal epithelial erosion), 2 (erosion with or without focal ulceration), or 3 (extended ulceration with or without granulation tissue). Mucosal bleeding was graded as 0 (absent), 1 (sporadic), 2 (diffuse within the lamina propria with or without patchy distribution), or 3 (diffuse with multiple affected areas). Lesion area was defined as the percentage of mucosal surface involved by erosions or ulcerations and scored as 0 (0%), 1 (1–25%), 2 (26–50%), or 3 (51–100%). Individual parameter scores were summed to generate a cumulative score (0–12), corresponding to normal (0–2), mild (3–5), moderate (6–9), or severe (10–12) inflammation. Scoring was performed in a blinded manner according to established criteria for DSS-induced colitis in mice.

### Fecal sample collection

At least three fresh fecal pellets were collected from mice either by directly placing them into a sterile microtube using standard scuffing techniques approved by the UBC Animal Care Committee or by using clean forceps to pick them up from the surface of a clean cage immediately after defecation.

### Mucus isolation

Isolation of fecal-associated mucus was performed on approximately 5 g of either frozen or fresh mouse pellets. The feces were placed directly into a 2 ml tube containing 1 ml of mucus extraction buffer (6 M guanidine hydrochloride (GuCl), 0.1 M Tris, 5 mM EDTA in filter-sterilized water with 10X cOmplete Protease Inhibitors. The solution was homogenized using a mechanical homogenizer (25 Hz for 5 minutes). All samples were then extracted overnight at 4°C on a rotator. The samples were pelleted by centrifugation (23,000 xg, 4°C, for 20 minutes), and the pellets were resuspended and re-extracted three more times with an equal amount of extraction buffer for 4 hours at 4°C, with the concentration of protease inhibitors decreasing by half for each round. Following these extraction steps and removal of the supernatant, samples were reduced twice with 100 mM dithiothreitol (DTT) suspended in extraction buffer to solubilize the mucin (the first overnight at 37°C, and the second for 6 hours at 37°C). Soluble mucins were then alkylated with 300 mM iodoacetamide on a rotator O/N at RT in the dark. The reduced and alkylated mucins were dialyzed into dH2O using Repligen 100 kDa MWCO Dialysis Device or tubing. Following dialysis, samples were treated with Pierce High-Capacity Endotoxin Removal Resin in batches (1 ml of resin to 1–10 ml of mucins) packed into 10 ml Pierce Centrifuge Columns. The dialyzed material was then freeze-dried overnight and stored at −20°C until further analysis via sialylomics and *O*-glycomics described below.

### Sialylomics analysis by ultra-high-performance liquid chromatography–mass spectrometry (uHPLC–MS)

The methods and materials used in this experiment are detailed in a previous publication^94^. In brief, a stock solution of 100 µg/mL sialic acid mixture (SA mix) in ultrapure water—containing 100 µg/mL each of Neu5Ac, Neu5Gc, 9-*O*-Neu5Ac₂, 4-*O*-Neu5Ac₂, Kdn, and Kdo—was prepared to create a range of calibrator solutions (0 to 25,000 ng/mL). Ten microliters of 10 µg/mL of ¹³C₃-Neu5Ac in ultrapure H₂O was added to each 100 µL aliquot of the calibrator solutions and samples (each containing 200 µg of mucus protein mass). These samples and calibrator solutions were then dried for total Sia analysis. For hydrolysis, 200 µL of 2 M acetic acid (AcOH) was added to the dried samples, followed by vortex mixing and incubation at 80 °C for 2 hours. The samples were then cooled to room temperature and labeled with 400 µL of a 24 mM 4,5-dimethylbenzene-1,2-diamine (DMBA) solution in 2 M AcOH at 60 °C for 1 hour. After cooling, samples were subjected to C18 solid-phase extraction. The 1 mL C18 cartridge (Discovery) was conditioned with 1 mL aqueous 80% acetonitrile (ACN)/0.1% trifluoroacetic acid (TFA) twice, followed by 1 mL of ddH₂O twice. The derivatized sialic acids were loaded, washed with 2 mL ddH₂O, and eluted with two sequential 0.5 mL aliquots of aqueous 50% ACN/0.1% TFA. The eluate was dried *in vacuo* at room temperature and reconstituted in 100 µL of 30% methanol/0.1% formic acid before being transferred to LC vial inserts for LC–MS analysis.

Ultra-high-performance liquid chromatography (uHPLC) was performed on an Agilent 1290 Infinity system equipped with a binary pump, autosampler, and column compartment. DMBA- derivatized sialic acids were analyzed using a Phenomenex Kinetex Biphenyl column (100 mm × 2.1 mm, 2.6 µm particle size, 100 Å pore size) at 40 °C. The injection volume was 7 µL for QTOF–MS. Mobile phases A and B were H₂O and MeOH, respectively, both containing 0.1% formic acid. The gradient elution program was: 0–4.2 min, 30–36.2% B; 4.2–8 min, 38% B; 8– 11 min, 50% B; 11–11.1 min, 90% B; 12.1–12.2 min, 30% B. Mass spectrometry was conducted on an Agilent Technologies 6530 QTOF mass spectrometer with an Agilent Jet Stream electrospray ionization (ESI) source, operating in positive ion mode. The source drying gas (N₂) temperature was 300 °C with a flow rate of 11 L/min; sheath gas (N₂) temperature was 350 °C with a flow rate of 11 L/min; nebulizer pressure was 40 psig; source nozzle voltage was 1000 V; and capillary voltage was 3500 V. Compounds were identified using Agilent MassHunter’s

*Find-By-Formula* algorithm. The QTOF–MS spectral acquisition rate was 2 spectra/s, and the QTOF–MS/MS spectral acquisition rate was 1 spectrum/s for precursor ions and 2 spectra/s for product ions. Sialic acid species were quantified based on the response ratios of their respective molecular ions against DMBA. Data acquisition and processing were performed using the MassHunter Workstation software suite (Agilent Technologies): Data Acquisition Workstation (v B.06.01), Qualitative Analysis (v B.07.00), and Quantitative Analysis (v B.07.00).

### Resorcinol Assay

Sia content was quantified using a resorcinol-based colorimetric assay following an adaptation of a procedure by Bhavanandan et al.^95^. In brief, samples and Neu5Ac standards (40 μL, in water) were hydrolyzed and oxidized by mixing with 10 μL sodium periodate (32 mM in 300 mM HCl) and incubating on ice for 1 h. Chromogenic complexes of the free, oxidized sialic acids were formed by addition of resorcinol reagent (resorcinol, 55 mM, copper (II) sulfate, 0.25 mM, dissolved in 6 M HCl) and incubation at 80 °C for 45 min. After cooling, 100 μL tert-butanol was immediately added to each sample and samples were incubated at 37 °C for 5 min. 250 μL aliquots of all samples and calibrators were transferred to 96-well plates and absorbance was then measured at 630 nm using a UV/Vis spectrophotometer. Sialic acid concentrations were determined by comparison to a Neu5Ac standard curve processed in parallel, and either normalized to the total protein concentration or the total carbohydrate content as assessed by the phenol-sulfuric acid assay (see below).

### Phenol Sulphate Assay

Total carbohydrate content of mucus samples was quantified using a microplate-based adaptation of the phenol–sulfuric acid assay with glucose as the calibration standard ^96^. For each reaction, 30 µL of each sample or glucose standard was dispensed into a flat-bottom 96-well polystyrene microplate. Concentrated sulfuric acid (150 µL) was added to each well, followed by addition of 30 µL of 5% (w/v) phenol in water. Plates were gently mixed and incubated at 90 °C for 5 min to allow chromophore development. After cooling to RT for 5 min, absorbance was measured at 490 nm using a microplate reader. Total glycan content was calculated from a glucose standard curve, and values were subsequently normalized to the corresponding Muc2 protein concentrations determined independently using a DC protein assay.

### Glycome analysis by Ultra-high-performance liquid chromatography-Mass spectrophotometry (uHPLC-MS)

Extracted mucus from mouse fecal pellet samples was lyophilized overnight using Labconco lyophilizer after which the glycans were released from serine/threonine residues of Muc2 using Carlson’s reductive β-elimination. This method utilized a solution containing 50 mM sodium hydroxide (NaOH) and 1 M sodium borohydride (NaBH4) in a total volume of 100 µL at a temperature of 45 °C with an incubation time of 16 h. Samples were cooled to 25 °C and neutralized by adding 10 µL aliquots of 2 M acetic acid; vortex-mixing was done after each addition until bubbling ceased. The samples were then diluted to 1 mL using ultrapure H₂O and dried *in vacuo* using a Savant SPD121P SpeedVac concentrator connected to a Savant RVT5105 refrigerated vapor trap (ThermoFisher Scientific) until the volume was reduced to approximately 50 µL; samples were then lyophilized until completely dry. The released glycans then desalted by solid phase extraction (SPE). SPE was performed using 250 mg Supelco ENVI-Carb graphitic carbon cartridges (Sigma-Millipore). The cartridges were first conditioned with 3 mL of 80% aqueous acetonitrile (ACN) containing 0.1% trifluoroacetic acid (TFA), followed by washing with 6 mL of ultrapure H₂O. Positive pressure was applied at the top of the tube for all SPE procedures. Crude glycan samples then were dissolved in 500 µL of ultrapure H₂O and loaded onto the cartridges. The sample tubes were then rinsed with 200 µl of ultrapure H₂O, the rinses also being applied to the SPE cartridges. The cartridges were washed with 3 mL of ultrapure H₂O, and reduced *O*-glycans were eluted using four sequential 550 µL aliquots of 50% aqueous ACN with 0.1% TFA. The pooled eluates were dried *in vacuo* to approximately 50 µL, transferred to 200 µL polypropylene HPLC vial inserts, and lyophilized. Prior to mass spectral analysis, samples were re-dissolved in 50 µL of H₂O.

Liquid chromatography was conducted using the Agilent 1290 Infinity system, with a binary pump, autosampler, and a temperature-controlled sample compartment (Agilent Technologies, Santa Clara, CA, USA). For mass spectrometry, an Agilent 6530 Quadrupole Time-of-Flight (QToF) system with jet stream electrospray ionization (ESI) was employed. Glycan separation was achieved using a Hypercarb™ HPLC column (Thermo-Fisher; 100 mm × 2.1 mm; particle size-3 µm). Details of the MS parameters, separation gradient, and solvents used in LC separation can be found in **Table 3 and Table 4**. High-resolution data were acquired at a mass setting of 3200 *m*/*z*. To prevent instrumental drift, a reference ion solution containing 10 μM purine (*m*/*z* 119.0360 for [M-H]^-^) and 2.0 μM HP-0921 (*m*/*z* 966.0007 and 1033.9881 for [M-H]^-^and [M+HCOO]^-^, respectively) in 95:5 ACN was added post-column at 800 μL/min using an Agilent 1260 Infinity II isocratic pump (with 1/16th of it directed to the MS and the remainder recycled). Full-scan spectra were collected at a rate of 2 Hz over a mass range of 100 - 3200 *m*/*z* and saved in profile format. HPLC-MS data were acquired and analyzed using MassHunter Workstation software (Agilent Technologies): Data Acquisition Workstation (v B.06.01, SP1) and Qualitative Analysis (v B.07.00, SP2). A database was created containing formulas for various combinations of *N*-acetylhexosamine (HexNAc), hexose (Hex), fucose (Fuc), *N*-acetylneuraminic acid (Neu5Ac, *i.e.* sialic acid), and sulfonate (SO3, *i.e.* a sulfated monosaccharide), ensuring each combination included at least one reduced HexNAc. The MassHunter’s Find-by-Formula algorithm in the Qualitative Analysis package was used to identify formula matches within a ± 10.00 ppm mass accuracy limit and, when applicable, a ± 0.350 min retention time window for matching peaks in other samples. Peak areas and retention times were processed using Microsoft Excel.

**Table 3.** Parameters for QToF-MS in negative electrospray ionization (ESI) mode.

| **Parameter (units)** | **Value** |
| --- | --- |
| Sheath gas flow(L/min) | 10 |
| Sheath gas temperature (°C) | 350 |
| Drying gas flow (L/min) | 10 |
| Drying gas temperature (°C) | 300 |
| Nozzle voltage (V) | 1000 |
| Nebulizer (psi) | 35 |
| Fragmentor voltage (V) | 175 |

**Table 4.** Gradient conditions used in uHPLC for separation of glycan.

| **Time (min)** | **Mobile phase B (%)** | **Flowrate (μL/min)** |
| --- | --- | --- |
| **0.0** | 0 | 250 |
| **5.0** | 5 | 250 |
| **15.0** | 20 | 250 |
| **20.0** | 40 | 250 |
| **25.0** | 80 | 250 |
| **27.0** | 80 | 250 |
| **27.1** | 5 | 250 |

\* Mobile phase A was 10 mM ammonium bicarbonate; mobile phase B was 10 mM ammonium bicarbonate in 80% (v/v) acetonitrile

#### Statistical analysis

Glycans were analyzed and visualized in RStudio (R version 4.3.1) using a custom R script, which quantified the relative abundances of different glycan groups (e.g., neutral, acidic, fucosylated, sialylated) to compare their distributions across genotypes. Relative abundances of individual glycans were analyzed using linear models with genotype as the main effect and sex as a covariate. Because male and female mice exhibited distinct baseline glycomic profiles and were not evenly represented across all glycan features, estimated marginal means (emmeans) were used to derive genotype comparisons adjusted for sex-related variability. This approach allowed genotype effects to be evaluated independently of sex imbalance, reducing the likelihood of spurious associations driven by unequal group composition. Multiple testing correction was performed using the Benjamini–Hochberg false discovery rate (FDR) procedure. In cases where no glycans survived FDR correction, glycans exhibiting unadjusted p < 0.05 together with biologically meaningful effect sizes were reported as exploratory, hypothesis-generating findings, and their sex-stratified distributions were examined to confirm consistency in directionality across sexes.

### Fecal DNA Extraction

Total genomic DNA was extracted from mouse fecal pellets using the QIAamp Fast DNA Stool Mini Kit (Qiagen) with modifications to enhance microbial cell lysis. Approximately 50–100 mg of fecal material per sample was suspended in ASL buffer and subjected to mechanical disruption using 0.1 mm zirconia/silica beads in a bead-beater at 6.0 m/s for 40 seconds. Following bead beating, samples were incubated at 95°C for 5 minutes to further promote lysis. DNA was eluted in 100 µL of nuclease-free water and stored at –20°C until analysis. DNA quality and concentration were assessed using a NanoDrop spectrophotometer and agarose gel electrophoresis.

### 16S rRNA library preparation and next generation sequencing

16S rRNA library preparation for individual samples at Gut4Health (RRID:SCR_023673) was prepared similar to the method described in deWolfe and Wright^97^. Briefly, the V4 region of the 16S rRNA gene was amplified with barcode primers containing the index sequences using a KAPA HiFi HotStart Real-time PCR Master Mix (Roche). PCR product amplification and concentration was monitored on a QuantStudio 3 Real-Time PCR system (Applied Biosystems). Amplicon libraries were then purified using AMPure XP Beads (Beckman), normalized based on concentration, and then pooled equally. Library concentrations were verified using a Qubit TM dsDNA high sensitivity assay kit (Invitrogen) and KAPA Library Quantification Kit (Roche) following manufacturer details. The purified pooled libraries were submitted to the Bioinformatics + Sequencing Consortium at UBC which verifies the DNA quality and quantity using an Agilent high sensitivity DNA kit (Agilent) on an Agilent 2100 Bioanalyzer. Sequencing was performed on the Illumina MiSeq TM v2 platform with 2 x 250 paired end-read chemistry. This work was supported by resources made available through the Gut4Health Microbiome Core Facility at the BC Children’s Hospital Research Institute (<u>RRID: SCR_023673</u>).

### 16S rRNA gene amplicon processing and analysis (Qiime2)

Paired-end 16S rRNA gene amplicon reads (V4 region; 515F/806R) were processed in QIIME2 ^98^. Raw FASTQ files were imported using a sample manifest in Phred33 format as `SampleData[PairedEndSequencesWithQuality]`. Primer and adapter sequences were removed using the QIIME 2 cutadapt plugin (`trim-paired`) with the following parameters: forward primer GTGYCAGCMGCCGCGGTAA, forward adapter ATTAGAWACCCBNGTAGTCC, reverse primer GGACTACNVGGGTWTCTAAT, reverse adapter TTACCGCGGCKGCTGRCAC, and `--p-no-indels`. Trimming performance was evaluated from the cutadapt report; 66,351 read pairs were processed and 100% passed filters (no reads were lost due to length or quality trimming). Demultiplexed read quality profiles were visualized using qiime demux summarize to guide downstream denoising/truncation decisions. Denoising, quality filtering, and amplicon sequence variant (ASV) inference were performed using the QIIME 2 DADA2 plugin (`denoise-paired`)^99^. Reads were trimmed from the 5′ end by 15 bp for both forward and reverse reads (`--p-trim-left-f 15`, `--p-trim-left-r 15`) and truncated to a length of 230 bp for both reads (`--p-trunc-len-f 230`, `--p-trunc-len-r 230`). DADA2 outputs included an ASV feature table, representative ASV sequences, and denoising statistics; these were summarized using `qiime feature-table summarize`, `qiime feature-table tabulate-seqs`, and `qiime metadata tabulate`, with sample metadata provided for per-sample read accounting and downstream grouping. Putative chimeric ASVs were identified using VSEARCH de novo chimera detection (`qiime vsearch uchime-denovo`) ^100^ applied to the DADA2 feature table and representative sequences. Chimera statistics were inspected (`qiime metadata tabulate`), and chimeric features were removed from both the feature table and representative sequences using `qiime feature-table filter-features` and `qiime feature-table filter-seqs` with `--p-exclude-ids`. The resulting non-chimeric feature table was re-summarized to confirm filtering outcomes. A phylogenetic tree for diversity analyses was constructed from non-chimeric representative sequences using `qiime phylogeny align-to-tree-mafft-fasttree`, which performs MAFFT multiple sequence alignment, masking of hypervariable positions, and FastTree inference of an unrooted tree followed by midpoint rooting. Alpha and beta diversity were calculated using `qiime diversity core-metrics-phylogenetic` with rarefaction to a sampling depth of 21,296 reads per sample, retaining 43/43 samples and 915,728 features at that depth. Core metrics outputs included standard alpha-diversity and beta-diversity distance measures, with metadata used for ordination and group comparisons. Data (Principal Co-ordinate analyses, Bray Curtis Distance Metrics) were visualized on with Qiime2View using Emporer. Taxonomic classification of ASVs was performed using a pre-trained Naive Bayes classifier targeting the V4 region (`classify-sklearn`; classifier: `2022.10.backbone.v4.nb.sklearn-1.4.2.qza`). Taxonomy and community composition were visualized with `qiime taxa barplot` using the non-chimeric feature table and sample metadata.

#### Microbiota compositional analyses

Alpha-diversity plots were generated in R with assistance from the Qiime2R package (https://github.com/jbisanz/qiime2R). Differential abundance analyses of 16S rRNA gene sequencing data were performed in R using compositional methods implemented in ANCOM-BC2^101^ and MaAsLin2^53^. Models included genotype as the primary predictor, sex as a covariate, and cage as a random intercept, where repeated housing could contribute to non-independence among samples. Prevalence and library size filtering were applied according to tool-specific recommendations. FDR-adjusted p-values were used to define statistically significant taxa. Taxa exhibiting consistent genotype-associated trends with moderate-to-large effect sizes and unadjusted p < 0.05, but not surviving FDR correction, were reported separately as trend-level associations, reflecting the exploratory nature of microbiota analyses and the conservative behavior of multiple-testing correction in high-dimensional datasets.

### Quantitative Microbiome Profiling (QMP) Workflow

#### Quantitative PCR (qPCR)

To obtain absolute bacterial abundances we followed Jian et al (2020)^51^, with some modifications.

#### Generation of Bacteroides thetaiotaomicron DNA Standard for Absolute Quantification

*Bacteroides thetaiotaomicron* (strain VPI-5482) was cultured in Brain Heart Infusion (BHI) medium supplemented with hemin under strict anaerobic conditions (85 % N₂, 10 % H₂, 5 % CO₂) at 37 °C. Bacteria cells were harvested by centrifugation and genomic DNA was extracted using the PureLink™ Genomic DNA Mini Kit (Invitrogen) following the manufacturer’s instructions. DNA yield and purity were verified spectrophotometrically (A₂₆₀/A₂₈₀ ratio ≈ 1.8–2.0), and aliquots were stored at –20 °C. Long-Product Whole-gene (LPW) PCR was employed to amplify a high-molecular-weight fragment of the *B. thetaiotaomicron* 16S rRNA genomic region for downstream usage as an absolute quantification standard. A master mix of total volume of 20 μL was prepared (7.5 μL nuclease-free water, 10 μL of 2X Master Mix, 1.5 μL primer mix (final concentration of 5 μM), and 20 ng of *B.theta* genomic DNA (final concentration 1 ng/µL) as template. The cycling protocol was as follows: The initial denaturation step was done at 95°C for 3 minutes. This was followed by 30 seconds at 95°C, 2nd denaturation step, annealing at 60°C for 30 seconds, and extension at 72°C for 40 seconds, for 35 cycles. A final extension at 72°C for 5 minutes was also performed. LPW primer sequences were: Forward (5′→3′) AGTTTGATCCTGGCTCAG:, Reverse (5′→3′): AGGCCCGGGAACGTATTCAC ^102^. PCR products were run on a 1 % agarose gel alongside a 1 kb Plus DNA ladder (Invitrogen Tra kit). The single expected band was excised and purified using a silica-membrane DNA extraction kit from gel to remove residual genomic DNA, primers, and nucleotides. DNA concentration was then determined using the Quant-iT™ PicoGreen™ dsDNA Assay Kit (Thermo Fisher Scientific) and expressed as pg/ µL. LPW primer pairs were aligned against the *B. thetaiotaomicron* VPI-5482 genome using NCBI BLAST. Five distinct amplicons were predicted, and their individual lengths were used to calculate the average amplicon size. Assuming an average molecular weight of 660 Da per base pair, the total number of DNA molecules per µL was calculated as:

Number of DNA Copies/µL = [(DNA conc. in pg/µL) × (6.022 × 10²³)] / [(average amplicon length in bp) × 660 × 10¹²]

where 6.022 × 10²³ is Avogadro’s constant, and 10¹² accounts for conversion of pg to gram. Based on the above calculation, a stock solution containing 10⁷ copies/2.5 µL of the LPW PCR product was prepared and serially diluted (10⁷–10² copies per 2.5 µL) to prepare standards for qPCR calibration. Semi-nested qPCR was then performed in duplicate to ensure amplification specificity and equivalent reaction efficiency between the LPW DNA standard and fecal bacterial DNA samples. Universal bacterial 16S rRNA gene primers (341F: 5′-CCTACGGGNGGCWGCAG-3′; 785R: 5′-GACTACHVGGGTATCTAATCC-3′) were used to target a shorter, internal region within the LPW amplicon. Each 20 µL qPCR reaction contained 10 µL iTaq™ Universal SYBR® Green Supermix (Bio-Rad), 0.4 µM of each primer, and 1 µL of template DNA. Amplification was carried out on a Bio-Rad CFX96 Real-Time PCR Detection System using: 95 °C for 5 min, followed by 40 cycles of 95 °C for 15 s and 60 °C for 30 s, with a melt-curve analysis (65–95 °C, 0.5 °C increments, 5 s per step). No-template controls (NTCs) were included in each run. Each qPCR plate included both the LPW DNA standards and fecal DNA samples from individual mice.

#### Quantitative real-time PCR

Quantitative real-time PCR was performed to determine absolute bacterial loads using universal 16S rRNA gene primers (e.g., 341F: 5’-CCTACGGGNGGCWGCAG-3’; 785R: 5’-GACTACHVGGGTATCTAATCC-3’) Each reaction was performed in triplicate in a 20 µL volume containing 10 µL SYBR Green Master Mix (e.g., Bio-Rad iTaq™ Universal SYBR® Green Supermix), 0.4 µM of each primer, and 1 µL of extracted DNA. Amplification was conducted on a Bio-Rad CFX96 Real-Time PCR Detection System under the following thermal cycling conditions: 95°C for 5 min, followed by 40 cycles of 95°C for 15 s, 60°C for 30 s, and a final melt curve analysis to verify product specificity. Standard curves were generated using ten-fold serial dilutions of purified PCR products or genomic DNA from reference bacterial strains with known 16S rRNA gene copy numbers. Only standard curves with amplification efficiencies between 90–100% and R² > 0.95 were used for quantification. Absolute abundances (16S rRNA gene copies per gram of feces) were calculated by interpolating sample Cq values against standard curves and normalizing to the initial fecal mass. The molecular weight of the *Bacteroides thetaiotaomicron* 16S rRNA gene (average length (n = 5 sequences) = 1371 bp, 660 Da/bp) was used to convert DNA mass to moles, which were further converted to absolute copy numbers using Avogadro’s constant. Copy numbers were normalized to the input DNA amount to calculate 16S rRNA gene copy number per ng DNA. To scale sequencing-based relative abundances, we incorporated (i) fecal sample weights, (ii) DNA extraction volumes and concentrations, and (iii) predicted 16S rRNA gene copy number per taxon derived from PICRUSt2 (marker_predicted_and_nsti.tsv). For each sample, the total DNA extracted was calculated from the measured concentration and elution volume. The total number of 16S rRNA gene copies per sample was then computed as the product of extracted DNA and qPCR-based copy number per ng DNA. This was normalized to fecal input mass to yield total 16S rRNA gene copies per gram of feces. Absolute abundances of individual taxa were calculated by multiplying their sequencing-derived relative abundance by the total 16S rRNA gene copies per gram. These values were corrected for taxon-specific 16S rRNA gene copy number predictions, yielding an estimated bacterial cell count per taxon per gram of feces. The final output included taxonomic assignments, raw read counts, relative abundances, absolute non-copy-corrected abundances, and copy-corrected bacterial cell counts for each taxon in each sample.

#### Absolute bacterial abundance (QMP) analyses

Quantitative microbiota profiling (QMP) data were analyzed to assess differences in total bacterial cell counts per gram of sample between genotypes. Normality of residuals was assessed using Shapiro–Wilk tests and visual inspection of Q–Q plots. Depending on data distribution, comparisons were performed using either parametric t-tests or nonparametric Wilcoxon rank-sum tests, with analyses conducted being both sex-adjusted and sex-stratified to ensure robustness of comparing the impact of genotypes.

### Neuraminidase Assay

Fresh fecal pellets (three per animal) were collected from male and female WT and IEC *Casd1^-/-^*mice, weighed, and homogenized in 100 mM sodium acetate buffer (pH 7.4) to a final concentration of 3.2% (v/w). Homogenates were centrifuged (15,000 × g, 4°C, 10 min), and supernatants were filtered (45 µm) and divided into paired aliquots treated with or without the neuraminidase inhibitor N-acetyl-2,3-dehydro-2-Deoxyneuraminic Acid (DANA; 10 mM final) as a specificity control. Following a 20-min incubation at room temperature, neuraminidase activity was assessed using the fluorogenic substrate 4-methylumbelliferyl-N-acetylneuraminic acid (4-MU-NeuNAc; 0.1 mM final) in a black 96-well plate and incubated at 37°C for 15 min in the dark. Reactions were terminated with sodium carbonate buffer (500 mM, pH 10.5), serially diluted to 1:16, and analyzed in triplicate with sodium acetate buffer alone as a blank. Fluorescence from released 4-methylumbelliferone was measured using a Tecan Infinite 200Pro plate reader (Ex 360 nm, Em 440 nm). Data were visualized and analyzed using RStudio (R version 4.3.2, R Foundation for Statistical Computing, Vienna, Austria) to see how 4-MU fluorescence activity differed between WT and IEC *Casd1^-/-^* fecal materials.

### SCFA Analysis

To analyze short-chain fatty acids protocols by Estaki et al.^103^ were utilized. Briefly, ∼50 mg of stool was homogenized with isopropyl alcohol, containing 2-ethylbutyric acid at 0.01% (v/v) as an internal standard, at 30 Hz for 13 minutes using metal beads. Homogenates were centrifuged twice, and the cleared supernatant was injected into a Trace 1300 gas chromatograph equipped with a flame-ionization detector with an AI1310 autosampler (ThermoFisher Scientific) in spitless mode. Data were processed using Chromeleon 7 software. Half of the fecal material was freeze-dried to measure the dry weight, and measurements are expressed as micromoles per gram of dry weight. Data was analyzed and plotted within RStudio using R software (Version 4.4.4; R Foundation for Statistical Computing).

### Statistical Analysis

All statistical analyses were conducted in RStudio (R version 4.4.1). Unless otherwise noted, data are represented as mean ± SD. Generally, depending on normality as determined by the Shapiro-Wilk test, either Welch’s t-test or the Wilcoxon rank-sum test was used to compare two groups in analyses without covariates. When covariates were present, outcomes were analyzed using linear or mixed-effects models, depending on data format. The key experimental factor in all analyses was genotype (WT vs. IEC *Casd1*^-/-^), with sex as a covariate in all models due to known sex-dependent differences in glycosylation, mucus biology, and host-microbiota interactions, as well as disproportionate representation of sex in some datasets. Fixed effects included genotype, sex, and region as appropriate. For outcomes measured repeatedly per animal (e.g., multiple regions for histopathology analysis), linear mixed-effects models were fit with mouse ID as a random intercept to account for animal correlation. When assumptions of normality or homoscedasticity were violated, Aligned Rank Transform (ART) ANOVA or ANCOVA was used to test main effects and interactions while preserving the factorial structure of the experimental design. For datasets averaged at the mouse level without repeated measures (e.g., mucus thickness, QMP results, and others), standard linear models adjusting for sex were used. In all datasets, statistical significance was declared at FDR-adjusted p < 0.05 where multiple testing was conducted (e.g., compositional analyses, SCFA concentration, Sialylomics). Findings that did not achieve this criterion but showed consistent direction of effect, biologically significant effect size, and unadjusted p < 0.05 were explicitly designated as exploratory trends and interpreted with caution. This strategy enabled reporting of biologically consistent signals in a transparent manner without over-interpretation of underpowered tests. Other statistical methods for complex compositional datasets (e.g., differential abundance, diversity indices) are described in their respective sections above.

## Acknowledgments

We gratefully acknowledge the major support for funding for this work from the Crohn’s and Colitis Canada Grants-in-Aid of Research (862392), and Michael Smith Health Research BC Scholar Award (Bergstrom).

## Ethics approval and consent to participate

Approval for the human analyses of the present study was given by the UBC Clinical Research Ethics Board (H21-3534, H16-03300, and H22-02645). Written informed consent to participate was obtained from all participants. No information given in this present study can be used to identify any study participant. The human studies abide by the principles articulated in the Declaration of Helsinki.

## Data availability

Data from this study are either included in this report or can be requested directly from the corresponding author.

## Supporting Information

This article contains supporting information.

## Declaration of generative AI and AI-assisted technologies in the manuscript preparation process

During the preparation of this work the corresponding author (K. Bergstrom) used ChatGPT 5.1/2 and Grammarly in order to proofread the original text for spelling and grammar without changing wording, and summarize and streamline complex methodologies (multi-omics, and statistical methods) to aid in clarity of approaches used; and assistance with troubleshooting R code for data analysis. After using this tool/service, the corresponding author reviewed and edited the content as needed and takes full responsibility for the content of the published article.

---

**Figure S1.**
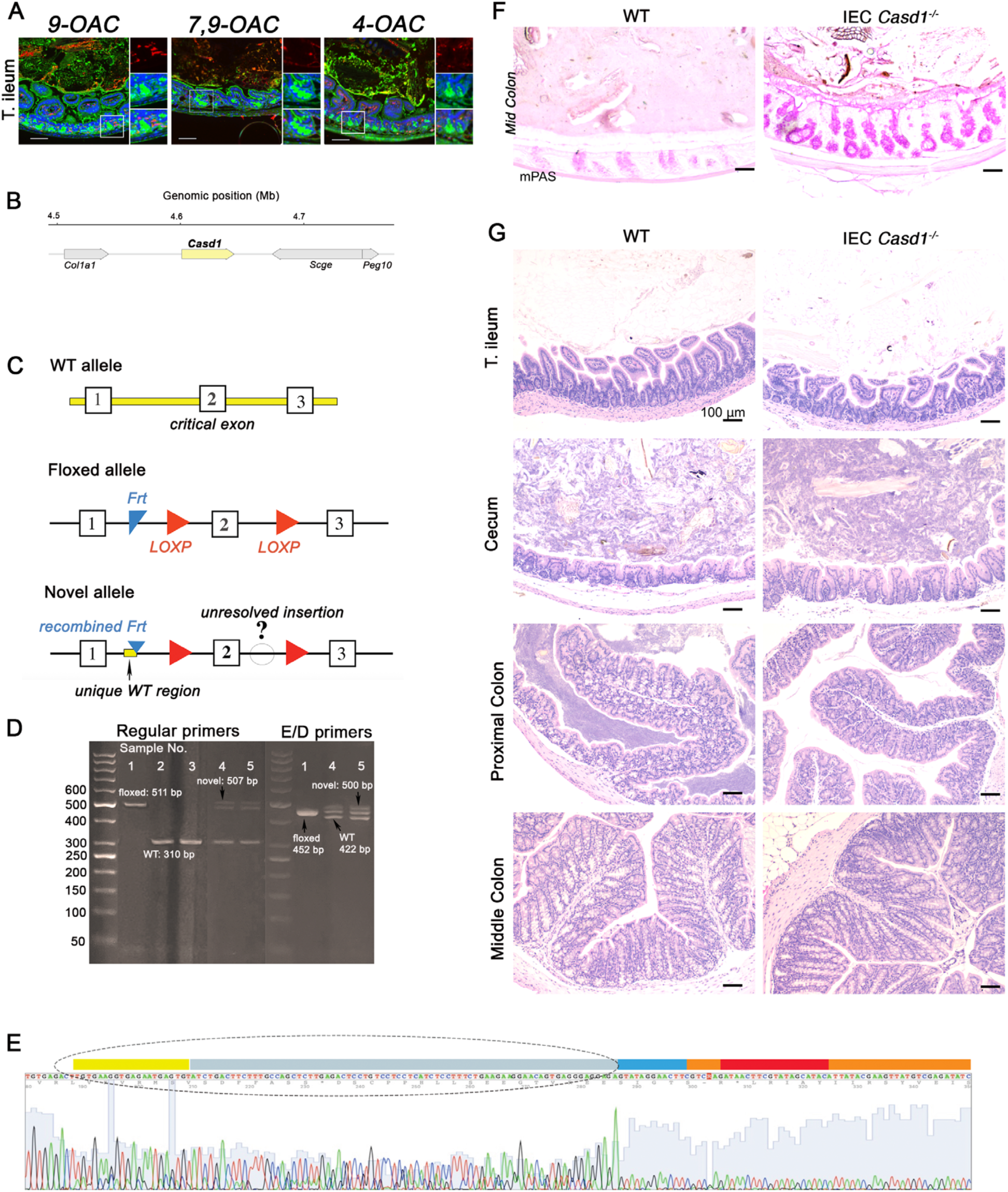
Loss of intestinal epithelial Casd1 does not alter tissue morphology in the intestinal tract. **A**. Epifluorescent imaging of *O*Ac-Sia variants on terminal ileal tissues **B**. Genomic neighborhood of Casd1 gene on mouse chromosome 6 (chr6). **C** Schematic representation of *Casd1* wild type (WT), floxed, and novel allele surronding the critical exon (exon 2). The floxed allele contains 2 LOXP sites flanking exon2 and a residual Frt sequence. The novel allele shows an unexpected pattern characterized by a partial retention of the Frt site, incorporation of a WT-unique sequence, and an unresolved inserted DNA region. **D**. PCR genotyping of *Casd1* alleles using regular & E/D primer sets amplifying the intronic region between exon 1 and 2, and the region spanning exon 2 and downstream LOXP site, respectively. PCR products were resolved on a 3% agarose gel with a 50-bp DNA ladder. **E**. Representative Sanger sequencing chromatogram of the unexpected PCR amplicon generated with the regular primers. The highlighted region indicates retention of a 99-bp WT-derived DNA sequence at the recombination junction. Color coding: blue: Frt, red: LOXP, yellow: WT unique sequences, orange: floxed unique sequences, gray: common areas between WT and floxed alleles. **F**. Representative mPAS staining of CFPE tissues. **G**. Representative H&E staining of tissues along the GI tract.

**Figure S2.**
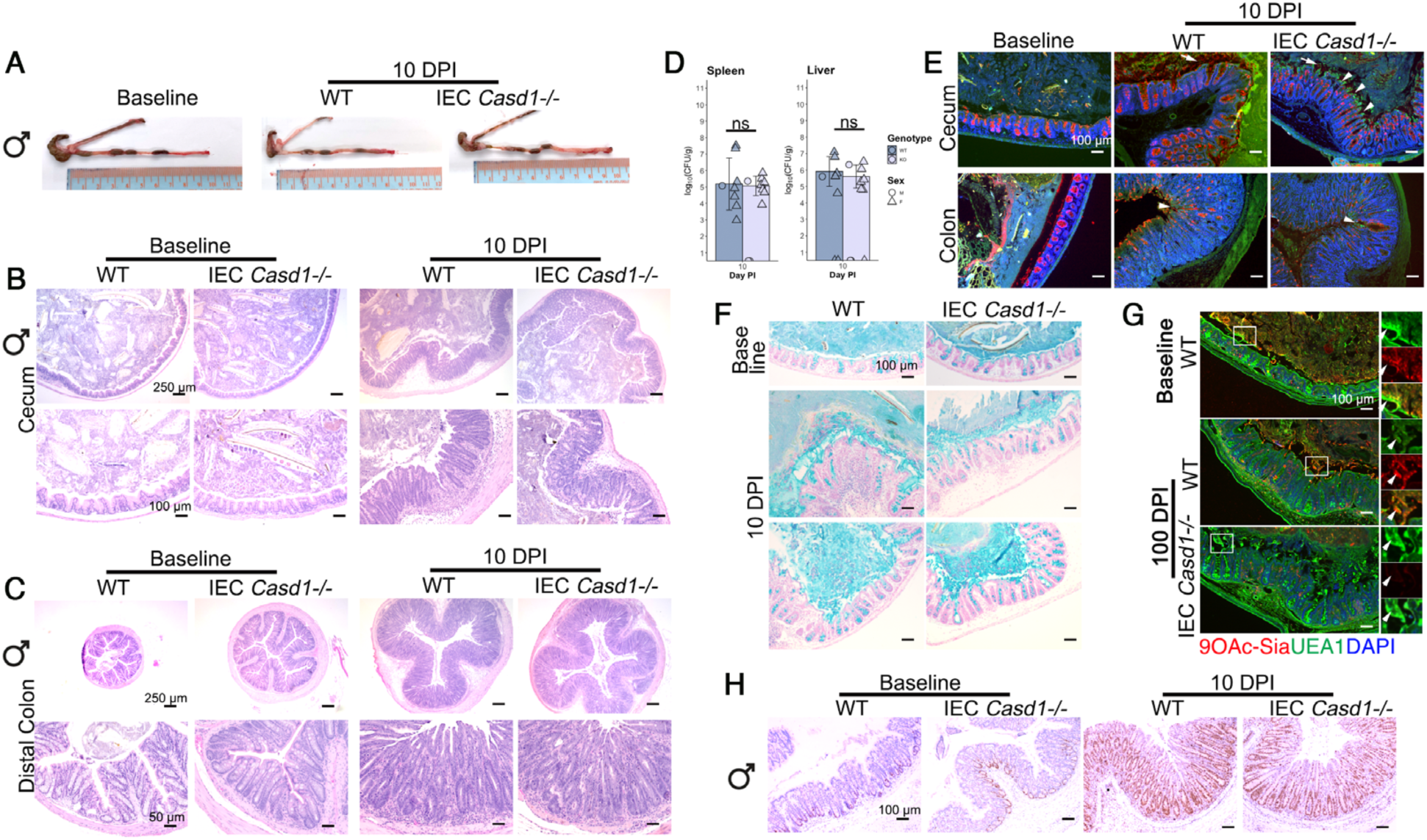
Assessment of colonization and pathology in male mice and cecal mucus responses. **A**. Picture of large intestine with terminal ileum. **B**. Representative low mag (upper panel) and high mag (lower panel) H&E staining of FFPE cecal tissues. **C**. Representative low mag (upper panel) and high mag (lower panel) H&E staining of FFPE colon tissues. **D**. Barplot (mean ± SD) of luminal *C. rodentium* burdens in the spleen and liver at experimental endpoint (10 DPI), with points representing individual mice. **E**. Epifluorescent labeling of LPS (green) and WGA (red) on CFPE tissues. Insets are magnified images of the corresponding boxed region. Arrows show secreted mucus. Arrowheads show mucosa-associated LPS typical of *C. rodentium* infection. **F**. Alcian Blue staining on CFPE cecal tissues. **G**. Epifluorescent labeling of 9-*O*Ac-Sia (red) and UEA 1 (green) on CFPE tissues. Insets are magnified images of the corresponding boxed region. Arrows show secreted mucus. **H**. Representative IHC for Ki67 on FFPE distal colon sections at baseline and experimental endpoint.

**Figure S3.**
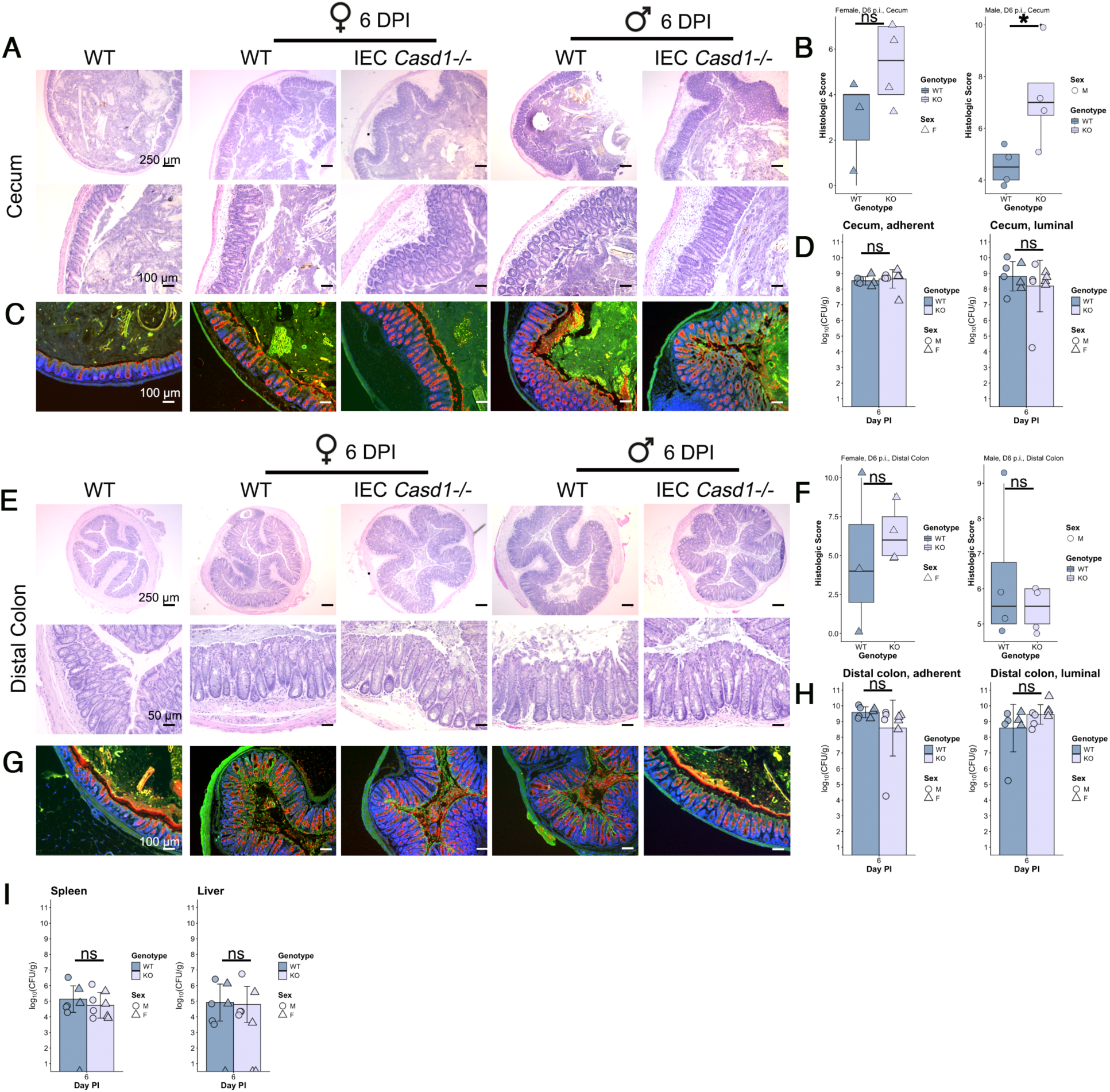
Pathology associated with *C. rodentium* at 6 DPI. **A**. Representative low mag (upper panel) and high mag (lower panel) H&E staining of FFPE cecal tissues. **B**. Boxplot of histologic damage scores in cecum at 6 DPI, with points representing individual mice (\**p* < 0.05; ns, *p* = 0.073; rank-based ANCOVA). **C**. Epifluorescent labeling of LPS (green) and WGA (red) on CFPE tissues. Insets are magnified images of the corresponding boxed region. Arrows show secreted mucus. Arrowheads show mucosa-associated LPS. **D**. Barplot (mean ± SD) of adherent vs. luminal *C. rodentium* burdens in the cecum at experimental endpoint (6 DPI), with points representing individual mice. **E**. Representative low mag (upper panel) and high mag (lower panel) H&E staining of FFPE colon tissues. **F**. Boxplot of histologic damage scores in distal colon at 6 DPI, with points representing individual mice (ns female, *p* = 0.558; ns male, *p* = 0.39; rank-based ANCOVA). **G**. Epifluorescent labeling of LPS (green) and WGA (red) on CFPE tissues. Insets are magnified images of the corresponding boxed region. Arrows show secreted mucus. Arrowheads show mucosa-associated LPS typical of *C. rodentium* infection. **H**. Barplot (mean ± SD) *C. rodentium* burdens in colon at experimental endpoint (6 DPI), with points representing individual mice. **I**. Barplot (mean ± SD) *C. rodentium* systemic burdens at experimental endpoint (6 DPI), with points representing individual mice.

